# Lab evolution, transcriptomics, and modeling reveal mechanisms of paraquat tolerance

**DOI:** 10.1101/2022.12.20.521246

**Authors:** Kevin Rychel, Justin Tan, Arjun Patel, Cameron Lamoureux, Ying Hefner, Richard Szubin, Josefin Johnsen, Elsayed Tharwat Tolba Mohamed, Patrick V. Phaneuf, Amitesh Anand, Connor A. Olson, Joon Ho Park, Anand V. Sastry, Laurence Yang, Adam M. Feist, Bernhard O. Palsson

## Abstract

Relationships between the genome, transcriptome, and metabolome underlie all evolved phenotypes. However, it has proved difficult to elucidate these relationships because of the high number of variables measured. A recently developed data analytic method for characterizing the transcriptome can simplify interpretation by grouping genes into independently modulated sets (iModulons). Here, we demonstrate how iModulons reveal deep understanding of the effects of causal mutations and metabolic rewiring. We use adaptive laboratory evolution to generate *E. coli* strains that tolerate high levels of the redox cycling compound paraquat, which produces reactive oxygen species (ROS). We combine resequencing, iModulons, and metabolic models to elucidate six interacting stress tolerance mechanisms: 1) modification of transport, 2) activation of ROS stress responses, 3) use of ROS-sensitive iron regulation, 4) motility, 5) broad transcriptional reallocation toward growth, and 6) metabolic rewiring to decrease NADH production. This work thus reveals the genome-scale systems biology of ROS tolerance.

**Graphical Abstract:** 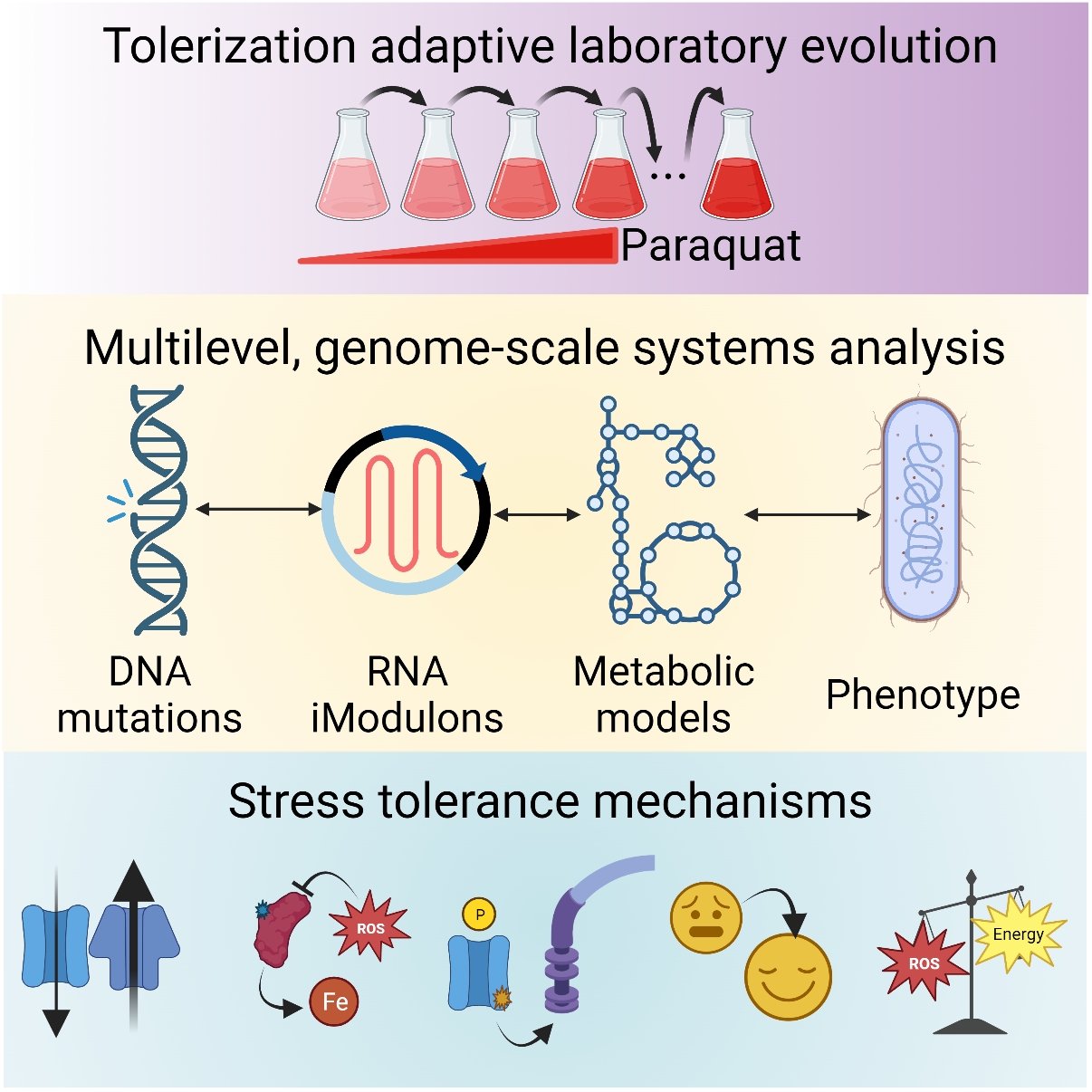

## Introduction

Omics technologies have enabled global understanding of cellular states at each level of the central dogma of biology. In particular, the falling cost of nucleotide sequencing has led to a dramatic increase in available genomic and transcriptomic datasets, allowing researchers to probe nucleotide changes in DNA and condition-dependent expression changes in RNA at unprecedented scale ^1^. With genome-scale metabolic models, we can also gain a global perspective on metabolic fluxes, and how they change based on genetic or expression perturbations ^2–4^. Each tool on its own has been successful in gaining novel biological insights, but an even deeper understanding can be achieved if they are made interoperable. Many approaches to integrate multiple omics data types are being developed ^5^, but the high number of variables and employment of complex “black-box” computational tools presents a problem for elucidating a clear, genome-scale understanding of biological systems across multiple levels of genomic, transcriptional, metabolic, and phenotypic changes.

Adaptive laboratory evolution (ALE) is an experimental procedure in which a microbial starting strain is grown in a selected condition for many generations, propagating when flasks reach a targeted density during repeated batch growth. This allows selection to enrich for mutant strains with improved fitness under the chosen condition ^6^. A tolerization ALE uses this procedure with increasing stressor concentrations, pushing cells to amplify stress tolerance mechanisms ^7^, thereby generating unique strains which are stress tolerance specialists. ALE strains are an excellent starting point for developing multi-omic approaches because they have a well-defined phenotype which arises from an average of only ~22 mutations ^8^. ALE mutations are highly informative for improving gene annotations, identifying fundamental biological principles and tradeoffs, designing bioproduction strains, and understanding antimicrobial resistance ^6,9^. However, it is difficult to interpret effects of mutations on regulators and enzymes without adding characterization from the transcriptome and metabolome.

The transcriptional regulatory network (TRN) employs transcription factors (TFs) which sense features of the cellular state and regulate the expression of genes in response. As transcriptomic data has been generated in rapidly growing numbers and deposited into online databases, it has become increasingly important to develop scalable methods which enable their interpretation. However, the typical method for transcriptional analysis, differentially expressed gene (DEG) analysis, is cumbersome for complex transcriptomic adjustments due to the high number of DEGs, and it does not easily capture the large-scale structure of the TRN. We seek to integrate signals from the TRN with mutations in the genome via biologically meaningful relationships, which is difficult if we do not first effectively decrease the number of transcriptomic variables.

A recently developed approach addresses this challenge by using independent component analysis (ICA) of large compendia of transcriptomic data to group genes into independently modulated sets (iModulons). The expression level (activity) of each group is computed in each sample, allowing systematic, large-scale analysis of the transcriptomic effect of adaptation to a new growth condition. Each iModulon is manually curated with predicted regulators and functions, bridging between the quantitative TRN and existing literature. iModulon activity levels can be used to infer the activity of their underlying regulators, and thus enable quantitative interrogation of the cell’s sensory systems. This approach has provided valuable insights into the TRNs of *Escherichia coli* ^10,11^ and several other organisms ^12–19^. iModulon analysis is supported by a developed codebase and online knowledgebase (iModulonDB.org) ^20,21^, which are publicly available. iModulons have already shown promise for analyzing transcriptional reallocation in tandem with mutations, which revealed important examples of the interplay between the genome and the transcriptome ^22–26^, but more work needs to be done to explain larger fractions of transcriptomic variance by systematically characterizing iModulon changes.

Downstream of the genome and gene expression, the state of the metabolic network is fundamental in determining cellular phenotypes. We have developed genome-scale metabolic and expression (ME) models, which compute optimal steady-state fluxes for all known reactions in a cell given mathematical constraints and an objective function ^2,27^. These models can be constrained with growth rates, uptake and secretion rates from metabolomic data, and transcriptomic data ^24,25^. Recent work has also incorporated the effects of biochemical stresses ^3,28,29^, enabling understanding of the cellular response to stress. Since ME models integrate phenotypic, metabolic, and transcriptional or proteomic data, they can be useful for supporting or refuting separate predictions made by analyzing genomic alterations.

The goal of the present study was to gain a genome-scale, multilevel, “white-box” understanding of a particular phenotype by leveraging ALE, genome sequencing, iModulons and ME modeling. Thus, we needed to select a well-defined phenotype of interest. We did so by employing ALE to generate *E. coli* strains which are specialized to tolerate a common herbicide, the redox-cycling compound paraquat (PQ). PQ is a redox-cycling compound, meaning that it can generate large amounts of reactive oxygen species (ROS) by stripping electrons from cellular electron carriers, such as NADH and NADPH, and reducing oxygen; this generates destructive superoxide ROS and regenerates the oxidizing agent to re-initiate the cycle ^30–32^. The ROS are particularly damaging to iron-containing enzymes and DNA. They decrease activity of important pathways, challenge the integrity of the genome, and inhibit growth ^3,32–36^.

Though the ROS response of *E. coli* is well understood and ROS are often delivered in the laboratory by PQ ^34^, some questions remain about how high levels of tolerance can be achieved: (i) In addition to the known proteins, which transporters and enzymes are involved in PQ cycling? (ii) What transcriptional alterations, specifically with respect to stress responses, metal homeostasis, and redox balance, are optimal? (iii) How can cells balance a tradeoff between generating NAD(P)H for energy and decreasing its production to prevent stress generation? Through our unique combination of systems biology techniques, we are able to shed new light on these questions. Their answers are informative for the fundamental biology of stress and metabolism, and for applications in pathology, antimicrobial design, and biomanufacturing.

This work provides a blueprint for combining ALE, mutational analysis, transcriptomics, computational biology, and phenotypic characterizations for stress-tolerant ALE strains, which emphasizes the rich insights provided by iModulon analysis. We begin by characterizing the strains and presenting an overview of the genomic and transcriptional changes. We then show that the effects of large DNA changes and TF mutations are easily quantified in the transcriptome. We also find an unexpected non-TF mutation that regulates motility regulons in our strains. Next, we disentangle the large fraction of the transcriptome which responds to changes in stress and growth phenotypes. Finally, we propose and model a metabolic mechanism for PQ tolerance which involves several interesting mutations and broad transcriptional reallocation. We show that the evolved strains employ a multi-pronged strategy of: (i) modifying membrane transport, (ii) using the SoxS and OxyR regulons to ensure stress readiness, (iii) allowing ROS-sensitive iron-sulfur (Fe-S) clusters to play a larger role in regulation of metal homeostasis, (iv) increasing motility, (v) shifting transcriptional allocation toward growth, and (vi) using fermentation to avert the PQ cycle. Taken together, these results elucidate a detailed, coherent, multilevel understanding of an important cellular phenotype by combining several cutting edge technologies in big data analytics and computational biology.

## Results

### Laboratory evolution increased tolerated PQ levels by 1000%

We evolved strains aerobically in minimal media with glucose under increasing PQ stress (**Figure 1A**). Our starting strain (0_0) was a derivative of *E. coli* K-12 MG1655 which had been pre-evolved to grow in minimal media with glucose ^37^. By using this media-adapted starting strain, the subsequent ALEs were enriched for mutations which improve stress tolerance, since the mutations that promote rapid growth under the culture conditions were already fixed. ALE was performed by steadily increasing PQ concentrations, first in three parallel first generation ALEs (1_0, 2_0, 3_0) and followed by eleven second generation ALEs (1_1, 1_2, …, 2_1, etc.) (**Figure 1A-B**). Parallelizing ALE replicates generated diverse strains and allowed for identification of common mutation targets which are more likely to be causal.

**Figure 1.**
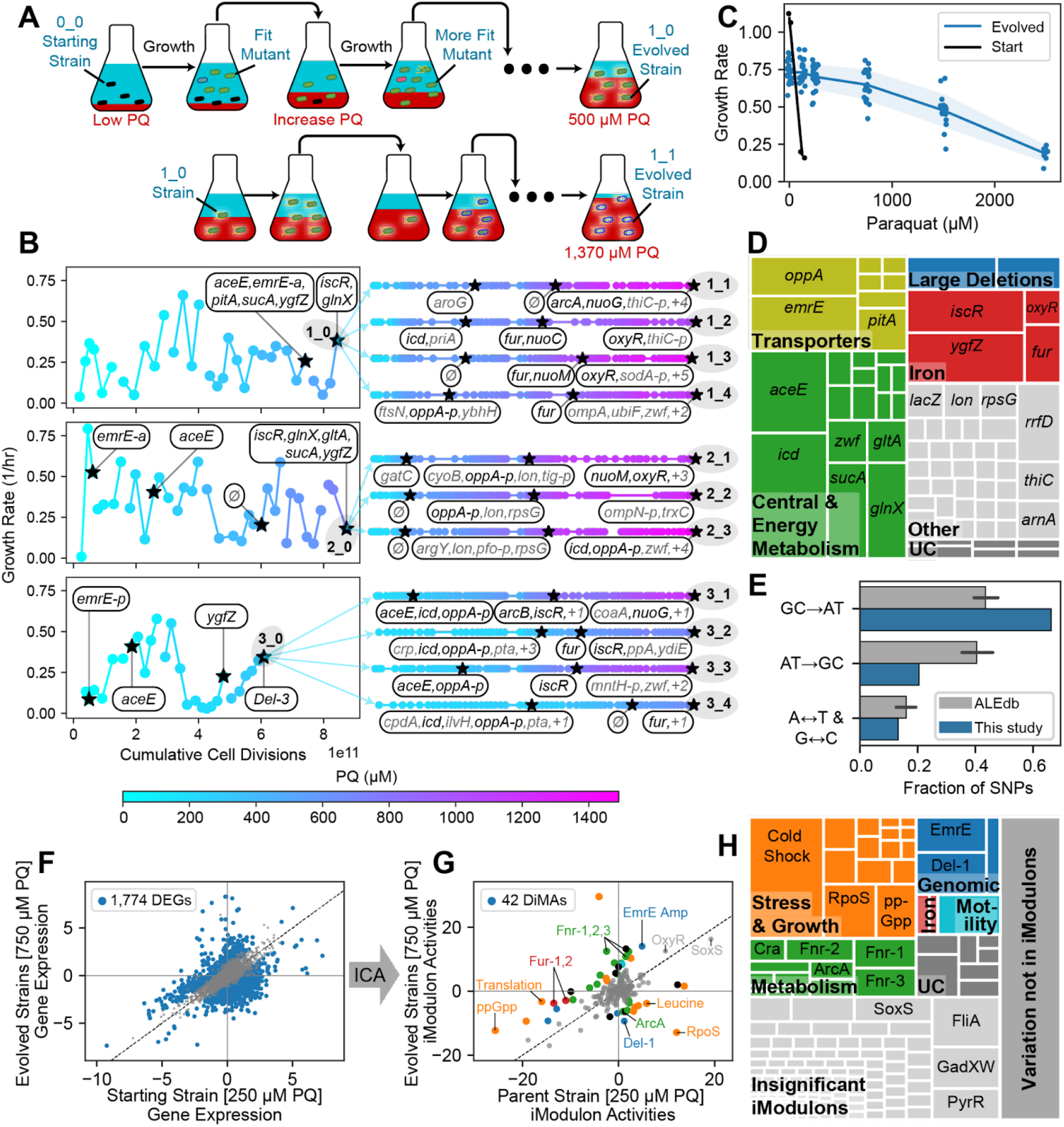
ALE increases PQ tolerance via changes to the genome and transcriptome. **(A)** Tolerization ALE process, showing mutant strains (cells with various appearances) in media with increasing stress concentrations (red). Example replicates are shown: 1_0 in the first generation and 1_1 in the second generation. **(B)** Points represent ALE flasks colored by their PQ concentration. The first generation of ALEs (strains 1_0, 2_0, and 3_0) are shown with each flask’s growth rate. ‘Cumulative cell divisions’ are estimated from the growth rate and time elapsed. Stars represent flasks that underwent DNA sequencing, and newly mutated genes are shown. Black colored genes are discussed in detail. **(C)** Growth rate for each strain at each PQ concentration. The starting strain cannot grow at 250 μM PQ, whereas some evolved strains reach up to 2500 μM PQ. Evolved strains grow slower than the starting strain in the absence of PQ. **(D)** Treemap of mutations in all strains, grouped by gene with intergenic mutations assigned to nearest genes. UC: Uncharacterized. See **Table S1**. **(E)** Fraction of SNP types in this study compared with all public ALE studies on ALEdb (aledb.org; mean ± 95% confidence interval). Each label corresponds to four of the twelve possible substitutions; for instance, “GC→AT” includes “G→A”, “G→T”, “C→A” and “C→T” substitutions. This study is enriched for mutations which decrease the GC content of the genome. **(F-G)** Comparison between the mean transcriptomes of the parent strain at 250 μM PQ vs. all evolved strains at 250 and 750 μM PQ. **(F)** DEG analysis, showing an intractably large number of DEGs. **(G)** Differential iModulon activity (DiMA) analysis, which compresses the differential transcriptomic changes into 42 DiMAs. DiMAs are colored by their category from panel (H). For more information about each iModulon, explore the PRECISE-1K E. coli dataset at iModulonDB.org and see **Table S2. (H)** Treemap of the explained variance of each iModulon in the transcriptome of the evolved strains. The map is first broken into three parts: the colorful region, composed of iModulons that are differentially activated after the evolution and categorized, the light gray region composed of iModulons that do not show a significant trend with evolution, and the dark gray region, representing the error in the iModulon decomposition.

After evolution, growth rates for each endpoint under different PQ concentrations were measured (**Figure 1C**). The starting strain’s growth was severely impaired by low PQ concentrations, with no growth at 250 μM PQ. The evolved strains showed a dramatic increase in the concentration of PQ they can tolerate while still growing; some endpoint strains tolerated 2500 μM. There was a fitness cost to the PQ tolerance, however: the strains no longer grew as well in the absence of PQ as the starting strain. This observation is consistent with the tradeoffs of the PQ tolerization mechanisms.

### Adaptive mutations reflect effects of PQ

Throughout the PQ ALE, a total of 222 mutations were observed, representing 111 unique sequence changes. Each mutation was assigned to its closest gene in the case of intergenic mutations, and 72 total genes were affected. Mutations were then categorized by their likely effects (**Figure 1D, Supplemental Table 1**). The largest category of mutated genes was central and energy metabolism-related (35%), which reflect the metabolic effects of PQ on redox balance. Transporters were also frequently mutated (16%), likely to prevent influx or promote efflux of PQ or other ROS. Iron and iron-sulfur (Fe-S) clusters are sensitive to oxidative stress ^34^, so we observed changes to iron regulators and Fe-S cluster synthesis genes (16%). Three large deletions, Del-1, Del-2, and E14 removal, were also notable (5%). Other mutations which were less convergent across endpoint strains (26%) were observed in ribosomal subunits, tRNAs, and *lon* protease, as well as across other parts of the metabolic network.

We performed DNA sequencing on several midpoint strains during the ALEs (**Figure 1B**), which provided insight into the most effective growth strategies since mutations tend to fix in the order of fitness benefit ^38^. We note that *emrE* and *aceE* are among the first genes to be affected in all three of our first generation strains.

An interesting pattern arose in the observed single nucleotide polymorphisms (SNPs): compared to other ALE projects available on ALEdb ^8^, they are highly enriched for changes from guanine or cytosine to adenine or thymine (**Figure 1E;** Fisher’s exact test p = 9.38*10^-5^). This enrichment was consistent with direct damage to DNA by ROS, since guanine is the most easily oxidized nucleotide ^33,39,40^. Thus, these mutations might not only improve cellular fitness through genomic and transcriptomic changes, but also by physically tolerizing cellular DNA to oxidation.

### iModulons enable analysis of complex transcriptomic changes

To identify transcriptomic adaptations, we performed RNAseq on the starting strain at 0 and 250 μM PQ, and on each evolved strain at 0, 250, and 750 μM PQ. In a comparison between the stressed samples for the pool of all evolved strains vs. the starting strain, we found 1,774 differentially expressed genes (DEGs) (**Figure 1F**), making detailed analysis using traditional transcriptomic methods challenging. Therefore, we applied iModulon analysis to enable interpretation.

The data was included in a large compendium of *E. coli* RNAseq data generated from a single wet lab protocol (PRECISE-1K ^11^). By leveraging over 1,000 samples across diverse conditions, this dataset facilitated machine learning of global transcriptomic patterns. Following our pipeline ^21^, we performed ICA on the full dataset. The result was a set of 201 iModulons, independently modulated gene sets which have similar expression patterns, along with their activities in each sample. Together, the iModulons constitute a quantitative regulatory structure which maps well to the known TRN, and can be used to reduce the dimensionality of the dataset. The set of PRECISE-1K iModulons was characterized in a separate study ^41^, and the iModulon structure, including interactive plots, search, and download functionality, is available at iModulonDB.org under *E. coli* PRECISE-1K ^20^.

iModulons enabled a global characterization of changes in the transcriptome. The evolved strains’ gene expression under PQ stress against the starting strain had only 42 statistically significant differential iModulon activities (DiMAs) (**Figure 1G**). These 42 iModulons made the analysis of the large-scale changes in the transcriptome tractable, and their observed activity changes could be related to the mutations fixed under ALE. We categorized the DiMAs and assigned mechanistic hypotheses which explain their changes (**Table S2**). Explained variance for all categories of significant and insignificant iModulons are shown in **Figure 1H**.

### A multilevel approach focused on explaining iModulon activities revealed the effects of mutations and phenotypes of evolved strains

Modifications to the genome can affect the transcriptome in several ways: large deletions and amplifications can directly alter the expression of genes involved, mutations in TFs can change the expression of their associated regulons, and the transcriptome can adjust due to changes in metabolites or other sensed processes that result from mutations. The latter type of alteration can be complicated by the fact that gene expression also regulates metabolite concentrations and sensed processes. In **Figure 2A**, we summarize how each of these types of relationships were observed in the evolved strains. iModulons play a central role in each highlighted mechanism, as evidenced by the full second column in **Figure 2A.** Their utility is a key outcome of this work. The combined analysis of genomic and transcriptional changes led us to six key cellular mechanisms of PQ tolerance (**Figure 2B**). Together, these mechanisms constitute a summary of the systems biology of PQ-generated ROS stress tolerance.

**Figure 2.**
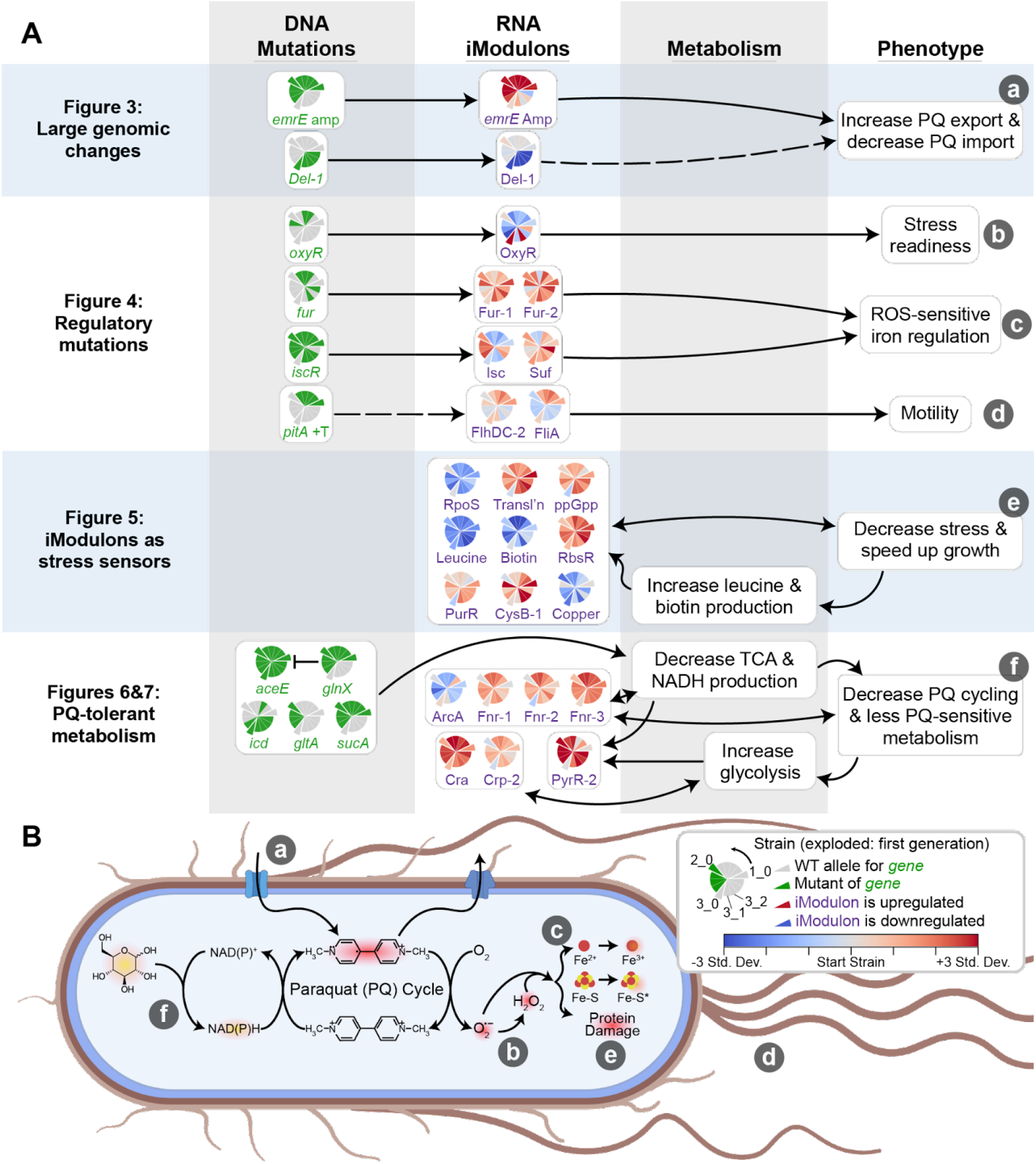
Multilevel approach reveals mechanisms of PQ tolerance. **(A)** Knowledge graph summarizing multilevel relationships between mutations, iModulons, metabolism, and phenotypes. Pie charts appearing in the two left columns indicate prevalence of given changes to the genome and transcriptome (legend in panel B), where wedges indicate strains. The protruding wedges correspond to the first generation of ALEs, with the wedges counterclockwise to them being their second generation descendants. For genes, green indicates the strain has mutations affecting it or its promoter. For iModulons, colors indicate the difference between the iModulon activity in the strain at 750 μM PQ and the starting strain at 250 μM PQ, normalized to the standard deviation of the iModulon activity across all of PRECISE-1K. Dashed lines represent relationships for which there is little existing literature. **(B)** Phenotypic changes target specific processes involved in PQ and ROS stress. Lowercase letters indicate elements from the rightmost column of (A). Entities which glow are reduced, and red indicates stress-related molecules.

### Large amplifications and deletions in the genome affect membrane transport

‘Genomic iModulons’ are transcriptomic modules which capture the effect of large changes to the genome, so they are of primary interest for obtaining genome-to-transcriptome relationships. In the PQ tolerant strains, the major genomic iModulons happen to all be associated with alterations in membrane transport.

The first mutation in each of the first-generation strains affected *emrE*, a multidrug efflux pump which pumps out PQ ^42^. In 1_0, 2_0, and their subsequent evolutions, genome coverage was increased approximately 42-fold in the region containing *emrE* (**Figure 3A**). This amplification was likely mediated by the flanking DLP12 prophage insertion sequence (IS) genes, specifically the IS3 transposase elements *insEF3* ^43^. ICA of the transcriptome recovered the amplified genes as an independent signal in the dataset, which we named the *emrE* Amp iModulon (called ROS TALE Amp-1 in PRECISE-1K ^11^ on iModulonDB.org ^20^). This iModulon showed elevated activity levels in all affected strains regardless of PQ concentration (**Figure 3C**). Thus, this case illustrates three levels in our multilevel approach (**Figure 2A**): it relates a clear mutational mechanism (transposase-mediated amplification) to a corresponding transcriptomic signal (*emrE* Amp iModulon) and beneficial phenotype (PQ efflux).

**Figure 3.**
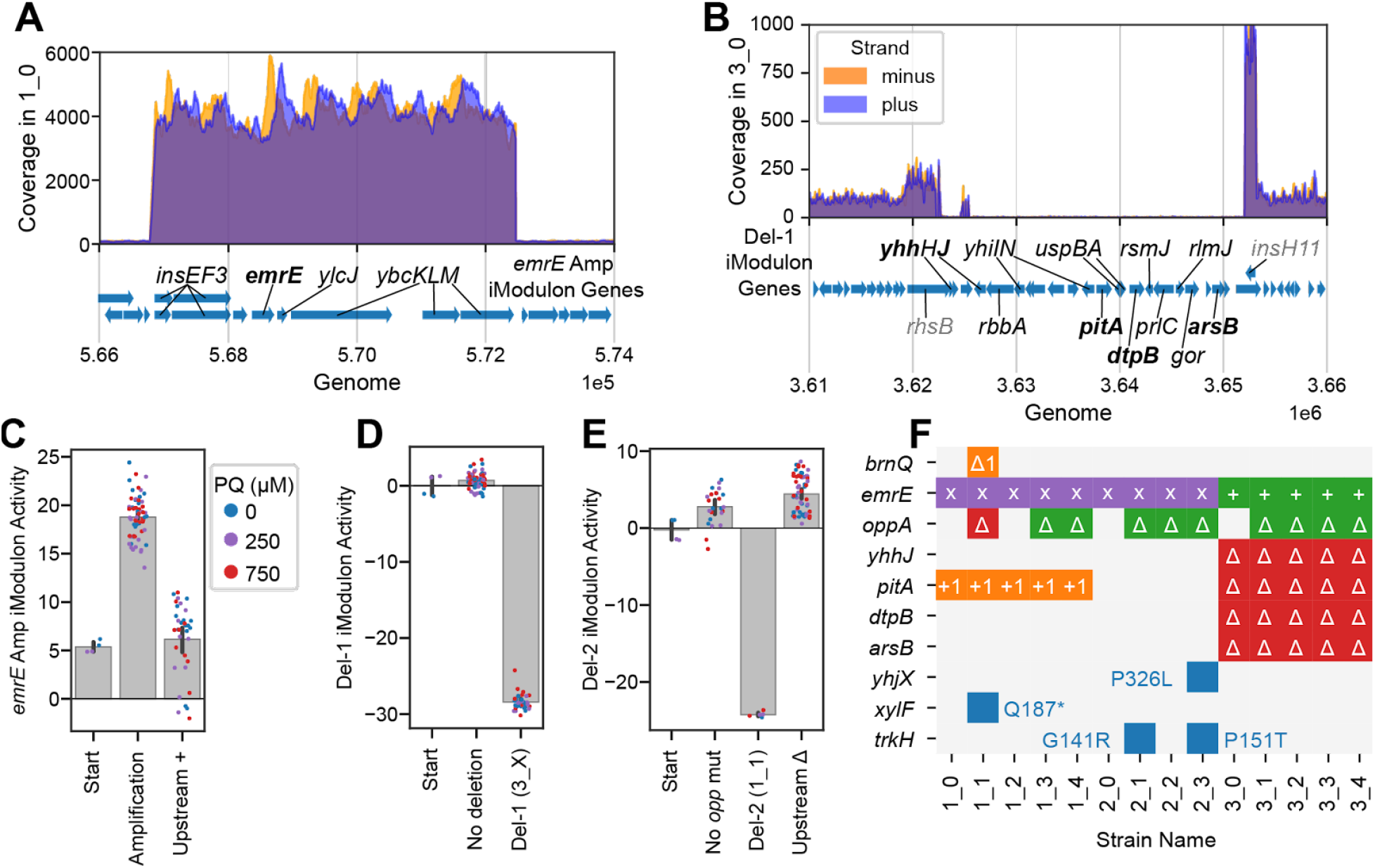
Consequences of deletions and amplifications affecting membrane transport are found in both genomes and transcriptomes. **(A)** Genome coverage in strain 1_0, which is representative of strains containing the emrE amplification, in the region of the amplification. Genes in the iModulon are labeled. **(B)** Genome coverage of strain 3_0 in the region of Del-1. Del-1 iModulon genes are shown in black, with flanking non-deleted, non-iModulon genes in gray, and transporters in bold. **(C-E)** iModulon activities for selected genomic iModulons. Bars indicate mean ± 95% confidence interval. Individual samples are color-coded by PQ concentration. Upstream + and Δ indicate insertions and deletions, respectively. **(F)** Color-coded table showing all observed mutations related to transporter genes. Purple x: amplification; green: upstream insertion (+) or deletion (Δ); blue: indicated SNP; orange: frameshift mutation within gene; red delta: complete gene deletion. The red area on the right indicates transporters deleted in the major 3_0 deletion.

We discuss Del-1, a large deletion that contains several transporters (**Figure 3B, D**), Del-2, a deletion of the *oppABCDF* operon (**Figure 3E**), and additional transporter mutations of potential interest (**Figure 3F**) in **Note S1.** We hypothesize that these mutations and their related genomic iModulons may have decreased influx of PQ or other oxidized molecules.

### Mutations in TFs alter the regulation of stress responses and iron homeostasis

‘Regulatory iModulons’ are iModulons which are statistically enriched with genes from a specific regulon, and their activity level quantifies the activity of the underlying TF. Thus, iModulon analysis reveals the effects of TF mutations in a convenient way.

The OxyR iModulon contains oxidative stress response genes, and its regulator, OxyR, responds to oxidative stress ^44^. Thus, we expected its activity level to correlate with PQ level. We found that for most strains, this is the case (p = 6.2*10^-5^). However, we observed three separate *oxyR* mutations which all fix OxyR iModulon activity levels at a level just below that of the stressed starting strain (**Figure 4A**), regardless of PQ concentration. We speculate that this level may be ideal because it enables quick detoxification of ROS, while higher levels would be proteomically expensive and/or induce growth-limiting levels of *oxyS* (which is regulated by OxyR and leads to growth arrest ^45^). Previous iModulon work in other ALEs found that fixing OxyR in the active conformation provided a fitness benefit^23^. Without the OxyR iModulon to quantify OxyR activity, it would have been much more difficult to define the effect of these mutations.

**Figure 4.**
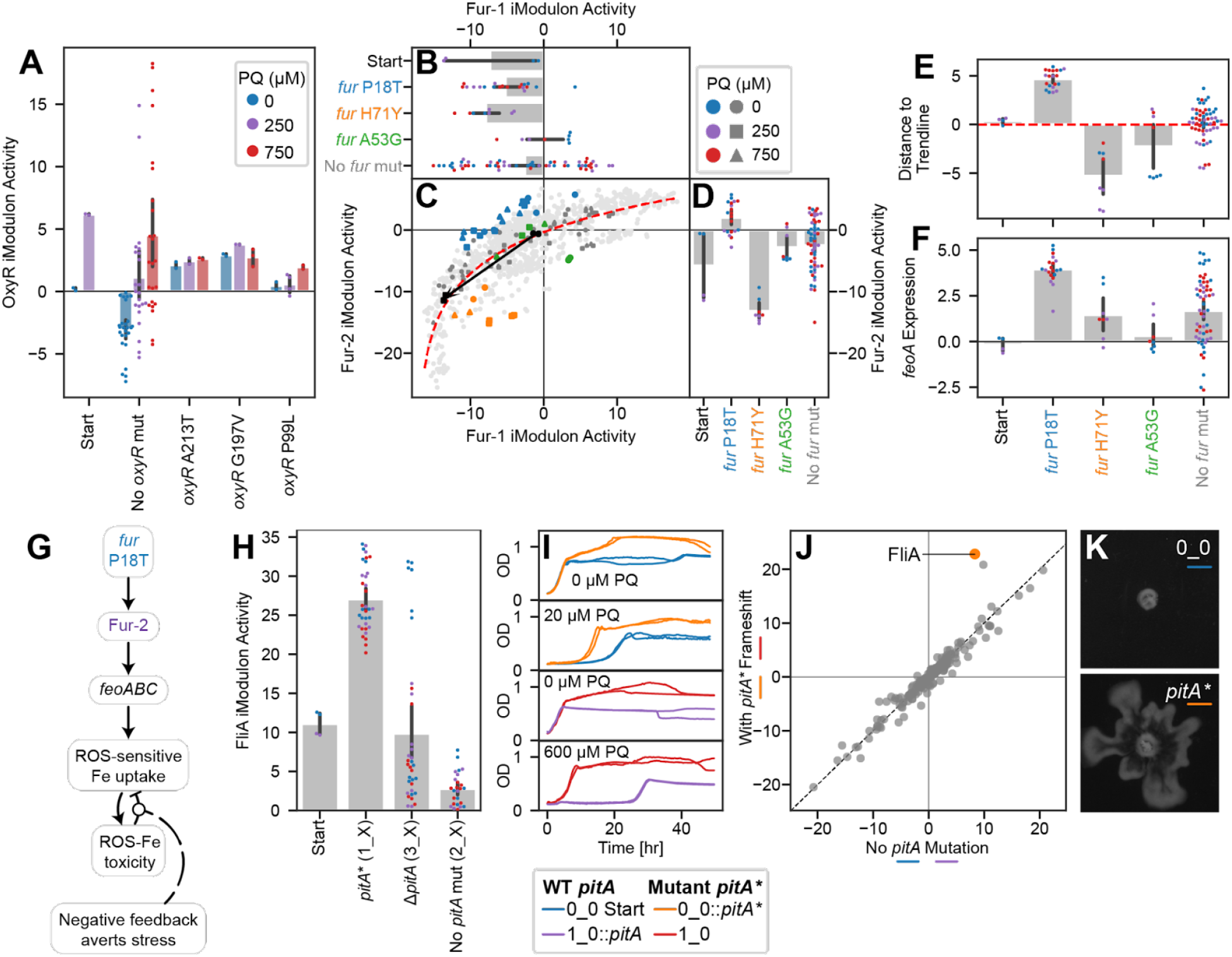
Mutations regulate stress response, iron metabolism, and motility iModulons in novel ways. Bars indicate mean ± 95% confidence interval. **(A)** OxyR iModulon activity is correlated with PQ in starting and evolved strains (Pearson R = 0.47, p = 6.2*10^-5^), except for the three strains which mutated *oxyR*. PQ colors in the legend also apply to panels (B, D, E-F, H). **(B-D)** Scatter plot of Fur-1 and Fur-2 iModulon activities with bar plots sharing axes. Light gray dots indicate other samples from PRECISE-1K. In (C), samples are colored by relevant mutations, and shapes indicate PQ concentrations according to the legends. A black arrow connects the starting strain samples between 0 and 250 μM PQ. In bar plots, point colors indicate PQ concentrations and label colors match with the scatter plots. The red trend line is a logarithmic curve fit to all samples in PRECISE-1k. Samples with the P18T mutation are above the trend line, indicating a preference for Fur-2. **(E)** Distances from each sample in this study to the trend line in (B), more clearly showing the preference for Fur-2 induced by P18T. **(F)** *feoA* expression, which is representative of the *feoABC* operon. Genes are upregulated by the *fur* P18T mutation. **(G)** Knowledge graph linking *fur* mutation to negative feedback which averts stress. **(H)** FliA iModulon activities by *pitA* mutation, showing an upregulation in the case of the frameshift *pitA* *, but not in the case of *pitA* deletion. (**I**) Growth curves for strains with and without the *pitA\** mutation as the only difference. The mutation contributes to higher final ODs under no stress, and shorter lag and faster growth under stress. (**J**) DiMA for strains 0_0 and 1_0 with and without the *pitA* frameshift mutation under PQ stress. Points indicate the mean of all relevant samples (individual conditions in duplicate; n=6 per axis). The strains with the mutation significantly activate FliA, one of the motility iModulons. The point near FliA is FlhDC-2, the other major motility iModulon. (**K**) Representative images of swarming in the 0_0 strain with (bottom) and without (top) the *pitA* * frameshift. Additional plots: **Figure S1;** Images for all swarming experiments: **Figure S2**.

Fur, the ferric uptake regulator, regulates two main iModulons (Fur-1 and Fur-2) whose activities have a nonlinear relationship which has been described previously ^46^ (**Figure 4B-D, S1A**). Three separate strains acquired *fur* P18T, which appears to shift Fur’s preference above the trend line, towards Fur-2 (**Figure 4E**). They specifically upregulate the expression of *feoABC*, a ROS-sensitive iron transporter ^47^ (**Figure 4F**). By using *feoABC* to strongly couple iron uptake to ROS levels, this mutation should prevent ROS-induced iron toxicity by preventing iron uptake under high stress (**Note S2; Figure 4G**). This mutation is of interest for further study, since it modifies the TRN in a unique way, and promotes a strategy of ROS-sensitive iron uptake that may be useful for production strains that are hampered by ROS.

The TFs IscR and SoxS also provide important insights. See **Figure S1** and **Note S3.**

These results highlight the efficacy of iModulon analysis for revealing TF mutational mechanisms. The Fur and IscR mechanisms predict that the ROS-sensitivity of Fe-S clusters can be used to couple iron uptake or utilization to ROS levels, which constitutes an interesting tolerance phenotype. The particular mutations ought to be introduced into production strains, where they could increase yields if ROS stresses are limiting. They can also be used to study TF-DNA interactions, since their effects on the transcriptome have already been elucidated here.

### An unexpected mutation in *pitA* upregulates motility

A frameshift in the phosphate transporter *pitA* led to a motile phenotype. This mutation occurred in 1_0 and its derivatives, and these strains also exhibited strong activation of motility-associated iModulons such as FliA (**Figure 4H**). There is no obvious connection between phosphate transport and motility, and the mutated strains were likely able to use the other phosphate transport system, *pstABCS*, to meet their phosphorus needs ^48^. Interestingly, the 3_0 strain deleted *pitA* as part of Del-1 (**Figure 3B**), and it did not exhibit the motility phenotype. Thus, to understand this mutation, we generated two new strains: *0_0::pitA\** and *1_0::pitA*, which added the mutation on its own to the starting strain and removed it in favor of the original *pitA* sequence in the evolved strain, respectively. We found that the mutation provided a growth advantage under PQ stress (**Figure 4I**). We also transcriptomically profiled the strains under the same conditions used for our other strains, and found that, particularly under PQ stress, the mutation exclusively perturbs the motility iModulons (**Figure 4J**). The change to the transcriptome was also reflected in the phenotype, as the mutant strains swarmed on agar plates while the wild-type *pitA* strains did not (**Figure 4K, S1**). The detailed mechanism of action linking the *pitA* mutant to motility remains to be elucidated.

An upregulation of anaerobic iModulons such as Fnr-3 in the ALE *pitA* mutants (**Figure S2A**) suggests a possible benefit for motility, in that it may be correlated with beneficial fermentation phenotypes discussed later, as has been previously studied ^49^ (**Note S4**).

This section illustrates the usefulness of our multilevel approach. After connecting mutations to their effects and predicting causes for DiMAs, we were left with an orphan mutation (*pitA*) and an unexplained DiMA (FliA). We predicted that the mutation caused the DiMA, and then we generated new strains to validate the prediction. The recapitulation of the expected iModulon change and swarming phenotype lends credibility to the iModulon method of elucidating mutational effects.

### Shifting from stress to growth explains activity of several iModulons

Regulatory iModulons can be used not only to understand the direct effects of mutations as described above, but also effects of changes to the processes that TFs sense. We have divided these types of changes in the PQ tolerant strains into two categories: those that respond to stress and growth (21% of the variance in the transcriptome; **Figure 1H**), and metabolic changes (10%). In this section we describe the former.

An important global tradeoff in the *E. coli* transcriptome is between growth and general stress readiness, which is governed by complex regulation ^50,51^. We previously identified a ‘fear-greed tradeoff’ between the RpoS and Translation iModulons, in which the activity levels of the two iModulons have a negative correlation; faster growing cells exhibit low RpoS and high Translation activity ^10,23,46,52,53^. The starting strain without stress is ‘greedy’, but it becomes ‘fearful’ upon addition of PQ, as expected (**Figure 5A-B**). The evolved strains, on the other hand, largely remain ‘greedy’ in the presence of PQ; they strongly downregulate RpoS (**Figure 5A**) and have higher translation activity than the stressed starting strain (**Figure 5B**). Translation activity is decreased relative to the starting strain in the absence of PQ, likely because of tradeoffs towards ROS stress readiness in the tolerized phenotype.

**Figure 5.**
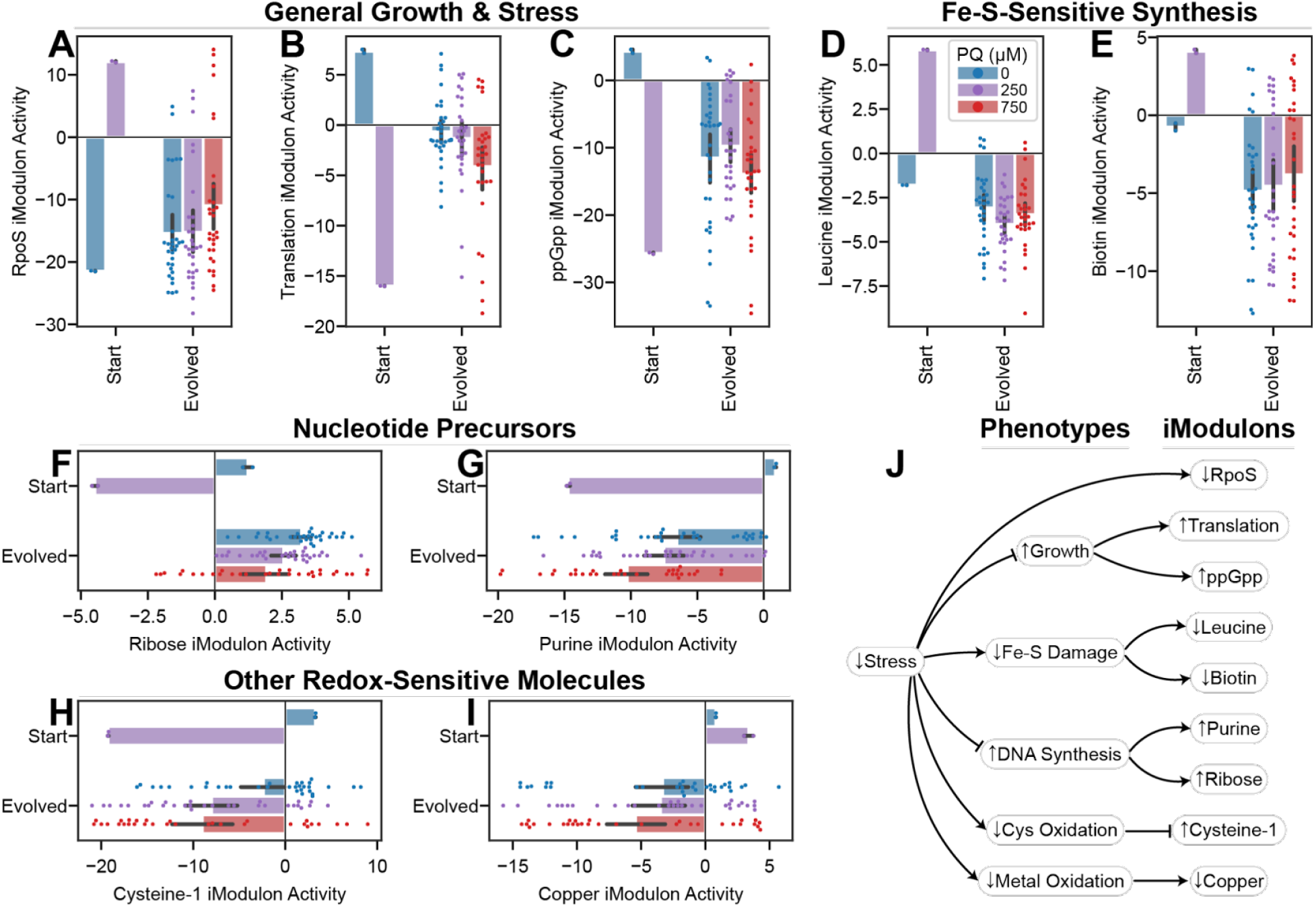
Changes to stress and growth explain the changes to activity in several iModulons. Mean iModulon activities ± 95% confidence interval; all plots use the legend in (D). P-values are false discovery rate corrected p-values from a comparison of stressed transcriptomes (250 and/or 750 μM PQ) between 0_0 and evolved strains. **(A)** RpoS activity, the general stress response, is downregulated (p = 0.017). **(B)** The Translation iModulon, ribosomes and translation machinery, is upregulated (p = 0.023). **(C)** The ppGpp iModulon, a large iModulon with many growth-related functions, follows a similar pattern to the Translation iModulon (p = 0.027). **(D)** The Leucine iModulon, which responds to leucine concentrations downstream of an Fe-S-dependent synthesis pathway, is downregulated after evolution, suggesting improved Fe-S metabolism (p = 0.0017). **(E)** The Biotin iModulon is downregulated after evolution. Biotin also depends on Fe-S-dependent synthesis (q = 0.017). **(F-I)** Ribose (p = 0.011), Purine (p = 0.036), Cysteine-1 (p = 0.025), and Copper (p = 0.034) iModulon activities behave differently in starting and evolved strains (**Note S5**). **(L)** Knowledge graph connecting decreased oxidative stress to each of the iModulon changes shown.

Despite the presence of stressors and the activation of specific ROS responses OxyR and SoxS, the global stress response is not activated in the evolved strains. There are two likely reasons for this: the stress signals are downregulated by the success of the evolved strategies of PQ tolerance, and the growth-inhibiting effects of RpoS have selected against strains with high RpoS activity. This work agrees with previous findings that ALE shifts allocation toward ‘greed’ ^10,23,52^. The decoupling of the ROS and general stress responses makes these strains ROS-response specialists, constituting a valuable adaptation strategy.

Two DiMAs reflect a decrease in oxidative damage by sensing Fe-S-dependent metabolites. The Leucine iModulon (**Figure 5D**) encodes the leucine biosynthesis pathway, which requires an Fe-S cluster and other metal-dependent enzymes that are sensitive to oxidative stress ^54^. Leucine feeds back to inhibit the iModulon’s expression ^55^. In the starting strain with PQ, oxidative damage likely leads to a decrease in leucine concentrations and an upregulation of the iModulon. By contrast, the evolved strains experience less stress, protect their Fe-S clusters, and therefore exhibit low Leucine iModulon activity. Similarly, the Biotin iModulon (**Figure 5E**) uses an Fe-S cluster in BioB to synthesize biotin ^56^, which then controls iModulon activity via regulation by BirA ^57^.

The activities of the ppGpp, Purine, Ribose, Cysteine-1, and Copper iModulons (**Figure 5C, F-I**) each also reflect decreased stress and a return to homeostasis in the evolved strains (**Figure 5J; Note S5**).

Thus, iModulons measure the entire sensory output of the TRN and allow us to mine the transcriptome for insights into many cellular processes. Because we also have an understanding of the stress phenotype of our cells, we predicted reasons for a large fraction of transcriptional alterations. This approach would be useful to any researcher seeking to enumerate phenotypic alterations in novel strains using only RNAseq data as a guide.

### Mutating central and energy metabolism genes decreases PQ cycling

We now turn to metabolism, which adds a fourth layer to our analysis and involves a complex interplay of effects from each level (**Figure 2A**). We show that enzyme mutations can suggest tolerance strategies, and then ME modeling can validate them. Finally, iModulon analysis can reveal how those strategies are organized and regulated by the cell.

The main metabolic mutations occur in the tricarboxylic acid (TCA) cycle. The second gene to mutate in all strains was *aceE* (**Figure 1B, D**). *aceE* encodes a subunit of pyruvate dehydrogenase (PDH), the entry point into the TCA cycle. *gltA, sucA*, and *icd* also mutate often, with *icd* being affected by e14 deletion and SNPs ^59^ (**Figure 6A**). These mutations would likely decrease the function of the enzymes, thus decreasing TCA cycle flux and production of NADH. These mutations suggest a tolerance benefit to decreasing NADH production. The likely reason for this benefit is that PQ uses electrons from NAD(P)H to reduce oxygen and generate stress ^60–62^. These mutations would decrease the available electrons to the PQ cycle and prevent stress generation. To decrease oxidative stress from PQ, the evolved strains perform less oxidative metabolism. The supplementary Fe-S and motility mechanisms (**Note S3, S4**) also shift strains away from NADH production.

**Figure 6.**
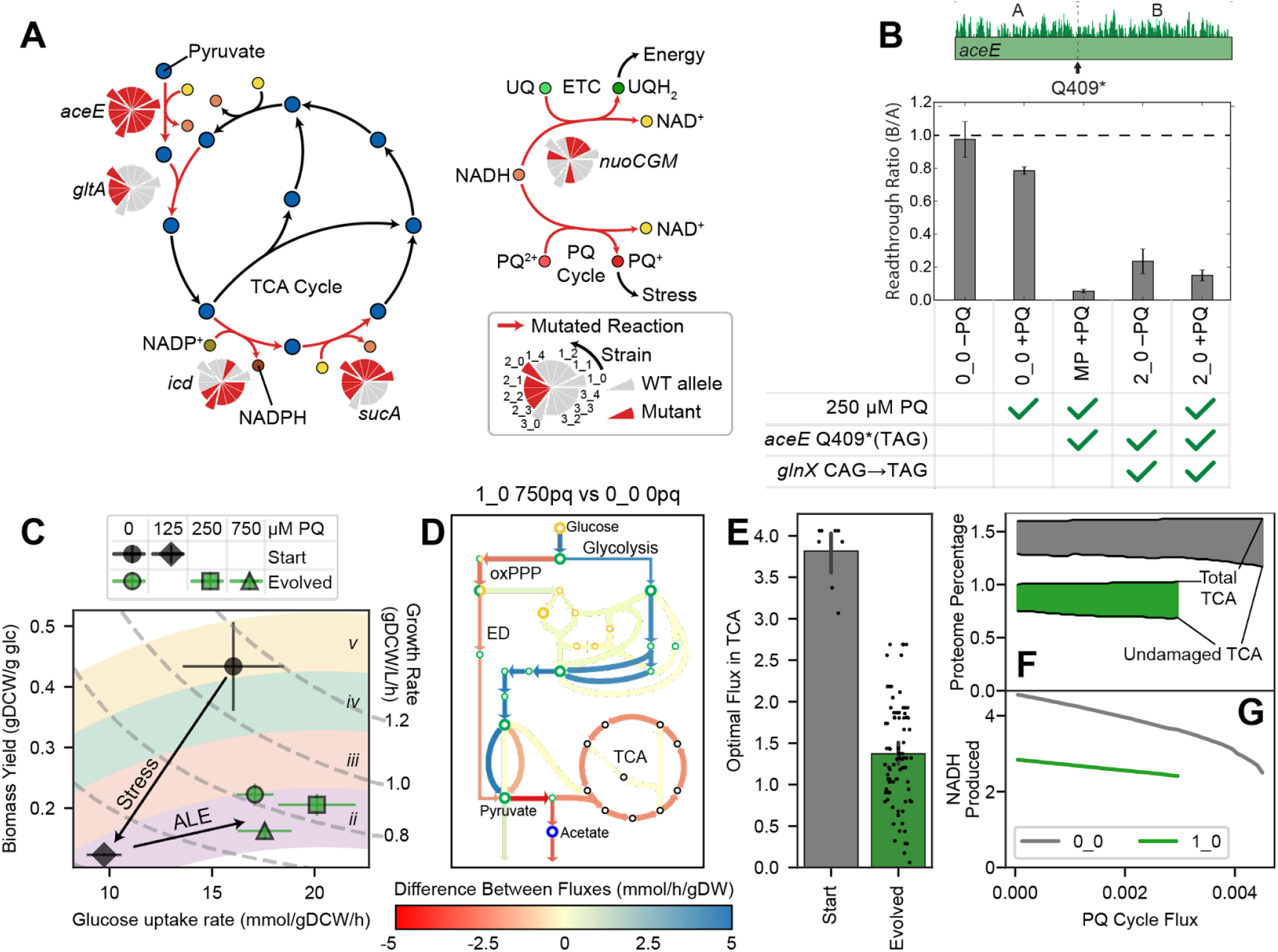
Mutations drive metabolic rerouting toward fermentation to avoid PQ cycling by decreasing NADH availability. (**A**) Simplified metabolic map of the TCA cycle and fate of NADH. Reactions catalyzed by mutated enzymes are shown in red and labeled with a pie chart indicating which strains have a wild-type (WT) or mutant allele. First generation strains in the pie chart protrude, with their descendants following them counter-clockwise. (**B**) Ribosome readthrough ratio in *aceE* from ribosome profiling, means ± standard deviation. The ratio B/A is the fraction of ribosomes bound downstream (B) vs. upstream (A) of the early amber stop codon (TAG) in *aceE*. The midpoint (MP) strain has *aceE* Q409* with WT *glnX*, whereas the 2_0 strain has both *aceE* Q409* and the *glnX* anticodon mutation that enables ribosomes to read through the amber stop codon. In evolved strains such as 2_0, PDH levels are decreased but not zero. **(C)** Aero-type plot ^58^ computed from measured growth rates and glucose uptake rates, where points represent means ± SEM, with constant growth rate isoclines. Colored regions labeled with roman numerals are aero-type regions as defined previously ^25^. Cells switch to a lower aero-type with PQ and increase their glucose uptake after evolution. **(D)** Flux differences from the OxidizeME model, comparing the starting strain with no PQ and a representative evolved strain at high PQ. Model was constrained by growth rate, glucose uptake rate, and RNAseq data (**Figure S4**). **(E)** Each point represents a TCA cycle reaction in the constrained OxidizeME models; models of evolved strains predict lower TCA cycle fluxes. **(F-G)** OxidizeME model results in mmol/gDCW/h for 0_0 and 1_0, constrained by growth rate, glucose uptake rate, and RNA expression. **(F)** As PQ cycle flux increases, the damaged fraction (filled in) of the TCA cycle increases. (**G**) NADH production decreases with PQ, but is more sensitive in 0_0. 0_0 can also carry more PQ cycle flux.

Loss of function (LOF) mutations in the TCA cycle come with a cost, since those pathways are the primary energy source for aerobic cells. Indeed, the evolved strains have decreased growth and translational activity under no stress relative to the starting strain, probably for this reason (**Figure 1C, 5B**). During ALE, the strains must therefore balance a tradeoff: generate enough NADH to grow and repair themselves, but not so much as to over-empower the PQ cycle. The tradeoff is embodied by an interesting interaction between mutations the *aceE* and *glnX* mutations, in which a tRNA mutation partially restored PDH activity (**Figure 6B; Note S6**).

We summarize all metabolic mutations in **Figure S3** and **Table S1**. We also observe mutations in enzymes involved in the utilization of NADH (**Figure S3A**; **Note S7**). Next, we characterize the strains using a variety of tools to test this explanation for the selection for TCA cycle mutations.

### Metabolic rewiring towards a lower aero-type decreases PQ sensitivity and flux in evolved strains

We quantified glucose uptake for each strain at various PQ levels (**Note S8**), and generated a plot comparing biomass yield per gram of glucose to the glucose uptake rate (**Figure 6C**). This rate-yield plane has been characterized in past studies ^25,58^, which revealed distinct energy generation strategies (aero-types) for each position in the plane. Samples with high biomass yields are in the highest aero-type (aero-type *v*), which represents efficient aerobic growth, whereas lower aero-types are associated with lower aerobicity and secretion of organic acids. The higher aero-types pump more protons across the inner membrane than the lower aerotypes^25^.

In **Figure 6C**, we observe a switch to a lower aero-type in the starting strain upon PQ exposure, since ROS damage decreases growth rate and particularly damages respiration. In the evolved strains, the lower aero-type is maintained even when no PQ is present. The aero-type change is likely due to the TCA cycle-related mutations, which we predicted would decrease respiration. However, the evolved samples also shift rightward, increasing their glucose uptake and total metabolic flux, enabling them to maintain growth under stress. Their position in the plane doesn’t vary much with PQ concentration, indicating decreased sensitivity.

To characterize metabolism *in silico*, we used OxidizeME, a genome-scale computational model of *E. coli* metabolism and expression (ME) which incorporates ROS stress effects ^3^. We constrained the model using each strain’s growth rate, glucose uptake rate, and RNA expression, then simulated optimal steady states (**Figure 6D, S4**). Though we did not attempt to simulate the effects of mutations on the reaction rates, the optimal flux distributions in the evolved strains showed decreases in TCA cycle flux (**Figure 6E**), consistent with the predicted effects of the mutations.

In the absence of experimental methods for directly measuring PQ cycle flux, we computationally assessed the consequences of PQ cycle flux by varying it for the starting strain and a representative evolved strain (**Figure 6F-G**). Though total proteomic allocation to the TCA cycle was constrained to match the RNA expression, ROS damage to the Fe-S clusters in *acnA, fumAB*, and *sdhABCD* led to decreasing functional proteome fractions (**Figure 6F**). The starting strain relied more heavily on the TCA cycle; this made it more sensitive to PQ, as evidenced by the steeper slope in NADH production (**Figure 6G**). The starting strain was also able to grow at higher PQ fluxes, which is inefficient and exacerbates stress. Thus, tolerization both decreases sensitivity to lower PQ fluxes and prevents a steady state with high PQ flux.

The genome-scale OxidizeME model integrates the individual cellular processes and RNA expression changes which adjust the phenotype, and it elucidates key systems level tolerization strategies. Its results match expectations from mutational analysis.

### iModulon activities shift tolerant strains towards anaerobic metabolism and glycolysis

Finally, we discuss iModulons which regulate the metabolic rerouting presented above. The cellular oxidation state is sensed and regulated by ArcA and Fnr ^63^, whose iModulons are differentially activated in the evolved strains (**Figure 7A-D**). Both TFs sense redox balance, which shifts towards reduction in the evolved strains due to the successful tolerization: ArcA represses when the electron transport chain is in a reduced state ^64^, whereas Fnr repression ceases when Fe-S clusters are intact^65^ (**Figure 7E**). These transcriptional changes shift from aerobic respiration genes toward anaerobic fermentation genes ^63^ (despite the aerobic ALE conditions). This strategy maintains a lower aero-type and decreases reliance on NADH. Thus, this mechanism reinforces the decreased reliance on the TCA cycle brought on by the mutations, ultimately slowing PQ cycling.

**Figure 7.**
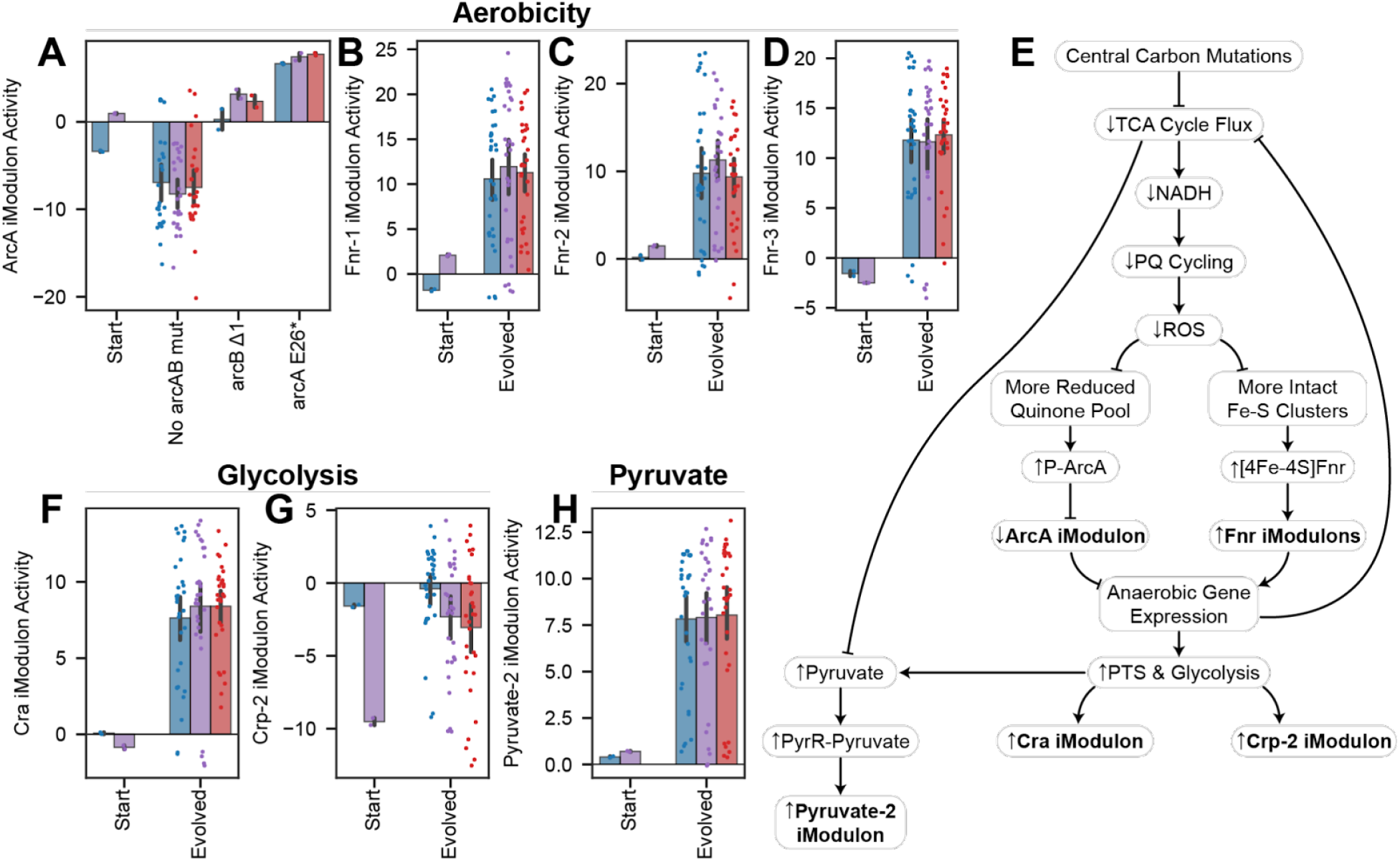
Mutations and iModulon reallocation drive metabolic rerouting toward fermentation to avoid PQ cycling. Bars indicate mean iModulon activities ±95% confidence interval. **(A)** ArcA iModulon activities are mostly decreased after evolution, except in the case of mutations to *arcAB* (p = 0.035). ArcA contains aerobic metabolism genes. **(B-D)** Fnr controls three iModulons with anaerobic metabolism genes, all of which are upregulated (p = 0.034, 0.030, 0.023). (**E**) Knowledge graph describing changes in the evolved strains connecting central carbon mutations to anaerobic and glycolytic gene expression, which decreases TCA cycle flux and ROS generation. **(F)** The Cra iModulon, which contains glycolytic genes that are repressed by Cra, is upregulated (p = 0.017). **(G)** The Crp-2 iModulon, which controls phosphotransferase systems, is upregulated (p = 0.022). **(H)** The Pyruvate-2 iModulon is upregulated (p = 0.012).

To meet energy needs with lower respiration, the cells increased their glycolytic activity, a change which is described by two DiMAs. Cra iModulon activity increases, indicating an increase in glycolytic flux (**Figure 7F**). Similarly, the Crp-2 iModulon returns to unstressed or intermediate levels in the evolved strains, which indicates a more active phosphotransfer system (**Figure 7G**). This transcriptomic change matches the rightward shift in the aero-type plot (**Figure 6C**). Finally, the LOF mutations downstream of pyruvate should increase pyruvate concentrations, which are sensed by the Pyruvate-2 iModulon and strongly upregulate it (**Figure 7H**). More details for all transcriptional mechanisms in this section are provided (**Note S9**).

In the past three sections, we showed that mutations and iModulon activity adjustments work together to enforce a low aero-type, PQ-tolerant metabolic network. The PQ tolerance stems from a decreased reliance on the TCA cycle and decreased NADH production, which leads to a metabolic network that supports less total PQ cycling and makes the system less sensitive to small amounts of PQ cycling. It is often difficult to interpret biological systems when genes, gene expression, and metabolic flux are all changing, but our multilevel interoperable approach using mutational analysis, iModulon activity changes, and genome-scale modeling produced a consistent and comprehensive interpretation of multiple data types.

## Discussion

In this study, we combined ALE with a detailed, systems-level transcriptomic analysis to comprehensively reveal mechanisms underlying PQ tolerance. The approach spanned four levels of analysis (**Figure 2A**): (i) genetic alterations and their predicted effects, (ii) transcriptomic adaptations along with up- and downstream inferences about their regulatory causes and physiological impact, (iii) metabolic fluxes calculated from genome-scale metabolic models, and (iv) phenotypic changes such as swarming motility. We found iModulon analysis of the transcriptome to be particularly revealing, as the TF activities could be readily quantified and utilized to infer a wealth of information about the phenotypic state. By combining these approaches into a coherent set of tolerization strategies, we presented a summary of the systems biology of ROS tolerance.

The evolved strains characterized herein achieved high tolerance through several mechanisms (**Figure 2B**). They promoted efflux of PQ via *emrE* segmental amplification, and precluded influx by mutating or deleting various other transporters. Inside the cells, PQ failed to generate as much ROS due to LOF mutations in and downregulation of NADH-producing pathways. To compensate for the decreased biomass yield of their metabolism, the cells increased glucose uptake and glycolytic flux. Since ROS interact with iron, some strains modified iron regulation via TF mutations that curtailed these systems when stress was high. These mutational and metabolic strategies led to a decrease in stress, which was sensed by the TRN and shifted various regulators toward faster growth.

The impact of this study is threefold. (i) We present biological insights of wide interest to researchers, including the growth/stress tradeoff of redox metabolism, the use of Fe-S clusters as a brake on iron uptake and metabolism, and novel interactions such as those between *pitA* and motility and between *aceE* and *glnX*. (ii) Acquired mutations and iModulon activities can become design variables for strain engineering, which frequently seeks to mitigate oxidative stress for bioproduction applications. (iii) We demonstrate an approach that utilizes iModulons to reveal a novel integrated perspective on adaptation to stress by understanding transcriptomic allocation.

Future studies should integrate additional data types into this framework. For instance, proteomics, endo-metabolomics, and chromatin immunoprecipitation of key TFs would be able to test various aspects of these hypotheses, better constrain models, and potentially uncover new insights. In addition, we encourage focused studies which characterize the mechanisms proposed here in greater detail.

Taken together, our results elucidate the systems biology of ROS tolerization using genome-scale datasets, computational models, and detailed literature review. Given the falling cost of RNAseq, development of laboratory evolution, and the availability of the pipeline developed here, we can expect that the systems biology of an increasing number of cellular functions and adaptations will be revealed.

## Supporting information

Table S1. Mutations in ROS tolerized strains

Table S2. Significantly differentially activated iModulons in ROS tolerized strains

## Acknowledgements

This work was funded by the Novo Nordisk Foundation Grant Numbers NNF10CC1016517 and NNF20CC0035580, and by National Institute of General Medical Sciences grant number GM057089. It used resources of the National Energy Research Scientific Computing Center, a DOE Office of Science User Facility supported by the Office of Science of the U.S. Department of Energy under Contract No. DE-AC02-05CH11231. We would like to thank Marc Abrams for his assistance with manuscript editing and Lachlan Munro for assistance with wet lab experiments. We would also like to thank Daniel Zielinski, Lei Yang, Emre Ozdemir, Tobias Alter, Ke Chen, and Jayanth Krishnan for helpful discussions. We thank Jason Yang for invaluable discussions and advice on an early draft of this manuscript. The graphical abstract was created in Biorender.

## Author contributions

K.R., J.T, C.A.O., J.H.P, L.Y., A.M.F, and B.O.P. designed the study. J.T., J.H.P, R.S., Y.H., A.A., C.A.O., J.J., and E.T.T.M performed experiments. K.R., J.T., A.V.S., P.V.P., and C.L. analyzed the data. A.P. performed simulations. K.R., J.T., and B.O.P. wrote the manuscript, with contributions from all the other co-authors.

## Declaration of interests

The authors declare no competing interests.

## STAR Methods

### Resource Availability

#### Lead contact

Further information and requests for resources and reagents should be directed to and will be fulfilled by the lead contact, Bernhard Palsson.

#### Materials availability

Strains generated in this study are available upon request.

#### Data and code availability

RNA-seq data have been deposited to GEO and are publicly available as of the date of publication, under accession numbers GSE134256 and GSE221314. DNA-seq data are available from aledb.org under the project “ROS”. iModulons and related data are available from iModulonDB.org under the dataset “*E. coli* PRECISE-1K”.

All original code and data to generate figures are available at github.com/SBRG/ROS-ALE, which also links to the alignment, ICA, and iModulon analysis workflows ^21^. It has been deposited at Zenodo and is publicly available as of the date of publication ^66^. The DOI is 10.5281/zenodo.7449004.

Any additional information required to reanalyze the data reported in this paper is available from the lead contact upon request.

### Experimental Model and Subject Details

#### Microbial strains

The starting strain (0_0) was an MG1655 K-12 *E. coli* strain which had been evolved for optimal growth on glucose as a carbon source in M9 minimal media ^37^. Mutations for the evolved strains are listed on aledb.org and in **Table S1.**

#### Culture conditions

Strains were grown overnight in M9 minimal media with 0.4% w/v glucose as a carbon source. Fresh media was inoculated with the overnight culture at an initial 600 nm optical density (OD) of 0.025. Cultures were aerated with a stir bar at 1100 rpm in a water bath maintained at 37°C until OD reached 0.5. 50 mM PQ was added to reach the desired concentration in stressed flasks. After 20 minutes, samples were harvested for transcriptomics or ribosome profiling.

### Method Details

#### Adaptive laboratory evolution

ALE was performed using a similar protocol to Mohamad *et al*. 2017 ^67^. Parallel cultures were started in M9 minimal medium by inoculation from isolated colonies. Evolution was performed in an automated platform with 15 mL working volume aerobic cultures maintained at 37°C and magnetically stirred at 1100 rpm. Growth was monitored by periodic measurement of the 600 nm OD on a Tecan Sunrise microplate reader, and cultures were passaged to fresh medium during exponential cell growth at an OD of approximately 0.3. Growth rates were determined for each batch by linear regression of ln(OD) versus time. At the time of passage, PQ concentration in the fresh medium batch was automatically increased if a growth rate of 0.08 h^-1^ had been met for 3 consecutive flasks. Samples were saved throughout the experiment by mixing equal parts culture and 50% v/v glycerol and storing at −80°C.

#### DNA sequencing and mutation calling

DNA was isolated as described ^68^. Total DNA was sampled from an overnight culture and immediately centrifuged for 5 min at 8,000 rpm. The supernatant was decanted, and the cell pellet was frozen at −80°C. Genomic DNA was isolated using a Quick-DNA Fungal/Bacterial Microprep Kit (Zymo Research) following the manufacturer’s protocol, including treatment with RNase A. Resequencing libraries were prepared using a Kapa Hyper Plus Kit (Roche Diagnostics) following the manufacturer’s protocol. Libraries were run on HiSeq and/or NextSeq (Illumina).

Sequencing reads were filtered and trimmed using AfterQC version 0.9.7 ^69^. We mapped reads to the *E. coli* K-12 MG1655 reference genome (NC_00913.3) using the breseq pipeline version 0.33.1 ^70^. Mutation analysis was performed using ALEdb ^8^.

#### Physiological characterization

Growth curves and exometabolomic samples were generated by inoculating cells from an overnight culture to a low OD using the same conditions as the ALE. For each strain, we started with 0 PQ. OD measurements and samples were taken at various time points until stationary phase was reached. We then passaged the cells into a new flask, stepped up the PQ concentration, and characterized the next curve, for concentrations 125, 250, 500, 750, 1500, and 2500 μM. We stopped if growth was not observed after 48 hours. For each flask, growth rates were determined by linear regression of ln(OD) versus time in the early exponential part of the curve.

We took cell culture samples at the same time as OD measurements for the starting strain at 0 and 125 μM PQ, and for the evolved strains at 0, 250, and 750 μM PQ. Samples were sterile filtered, and extracellular by-products were determined by high pressure liquid chromatography (HPLC). The filtrate was injected into an HPLC column (Aminex HPX-87H 125-0140). The concentrations of the detected compounds were determined by comparison to a normalized curve of known concentrations. Substrate uptake and secretion rates in the early exponential growth phase were calculated from the product of the growth rate and the slope from a linear regression of the grams (dry weight) (gDW) versus the substrate concentration. The biomass yield was calculated as the quotient of the growth rate and the glucose uptake rates during the exponential growth phase.

#### RNA Sequencing

3 mL of induced culture was added to 6 mL of RNAProtect Bacteria Reagent (Qiagen) and vortexed, then left at room temperature to incubate for 5 minutes. Cells were pelleted, resuspended in 400 μL elution buffer, and then split into two tubes with one kept as a spare. One pellet was then lysed enzymatically with addition of lysozyme, proteinase-K, and 20% SDS. SUPERase-In was added to maintain the integrity of the RNA. RNA isolation was then performed according to the RNeasy Mini Kit (Qiagen) protocol. rRNA was depleted using the Ribo-Zero rRNA Removal Kit for gram negative bacteria according to the protocol. Libraries were constructed for paired-end sequencing using a KAPA RNAseq Library Preparation kit. Reads were sequenced on the Illumina NextSeq platform.

As part of the PRECISE-1K dataset ^41^, transcriptomic reads were mapped using our pipeline (https://github.com/avsastry/modulome-workflow) ^21^ and run on Amazon Web Services Batch. First, raw read trimming was performed using Trim Galore with default options, followed by FastQC on the trimmed reads. Next, reads were aligned to the *E. coli* K-12 MG1655 reference genome (NC_000913.3) using Bowtie ^71^. The read direction was inferred using RSeQC ^72^. Read counts were generated using featureCounts ^73^. All quality control metrics were compiled using MultiQC ^74^ Finally, the expression dataset was reported in units of log-transformed transcripts per million (log(TPM)).

All included samples passed rigorous quality control, with “high-quality” defined as (i) passing the following FastQC checks: *per_base_sequence_quality, per_sequence_quality_scores, per_base_n_content, adaptor content;* (ii) having at least 500,000 reads mapped to the coding sequences of the reference genome (NC_000913.3); (iii) not being an outlier in a hierarchical clustering based on pairwise Pearson correlation between all samples in PRECISE-1K; and (iv) having a minimum Pearson correlation between biological replicates of 0.95.

#### Ribosome profiling

Ribosome profiling libraries were created using a modified version of the protocol outlined in Latif *et al*. ^75^. The protocol was modified to negate the effects of the addition of chloramphenicol by grinding frozen cells. 50 mL of cell culture was harvested by centrifugation for 4 minutes at 37°C in a 50 mL conical tube containing 0.4 g of sand. Supernatant was aspirated quickly and the pellet was flash frozen in liquid nitrogen. Pellets were transferred into a liquid nitrogen cooled mortar and pestle, 500 μL of lysis buffer was added, and the pellet was pulverized to lyse the cells. Lysate was transferred to a falcon tube to thaw on ice. The lysate was then centrifuged, and the supernatant was isolated to continue with the published protocol. Reads were sequenced on an Illumina HighSeq machine using a single end 50 bp kit.

Adaptors were removed from ribosome profiling reads using CutAdapt v1.8 ^76^, then mapped to the *E. coli* K-12 MG1655 reference genome (NC_000913.3) using bowtie ^71^. They were scored at the 3’ end to generate ribosome density profiles.

#### *Generation of* pitA *mutant strains*

The mutations referred to in **Figures 4H-K and S2** were introduced into the starting (0_0) and evolved (1_0) genomes using a Cas9-assisted Lambda Red homologous recombination method. Golden gate assembly was first used to construct a plasmid vector harboring both Cas9 and lambda red recombinase genes under the control of an L-arabinose inducible promoter, a single guide RNA sequence, and a donor fragment generated by PCR which contained the desired *pitA* +T mutation and around 200 bp flanking both sides of the Cas9 target cut site as directed by the guide RNA. After allowing cells harboring the plasmid to grow for 2 hours at 30°C, L-arabinose was added to the media and the cells were allowed to grow for 3 to 5 hours, at which time a portion of the culture was plated. Single colonies were screened using ARMS PCR. Amplicons spanning the mutation site, generated with primers annealing to the genome upstream and downstream of the sequence of the donor fragment contained in the plasmid, were confirmed with Sanger sequencing. Confirmed isolates were cured of the plasmid by growth at 37°C.

#### Cell motility assay

We performed motility assays in duplicate for each of the conditions shown in **Figure S2.** We mixed a tryptone broth (13 g tryptone and 7 g NaCl per liter of media) with 0.25% agar and the desired PQ level. We autoclaved the broths, then poured 25 mL into petri dishes and solidified them at room temperature overnight. Fresh colonies were spotted in the middle of the semi-solid agar with a toothpick. The plates were then incubated at 37°C for 6-8 hours and imaged on a Gel Imaging System.

### Quantification and statistical analysis

#### iModulon computation and curation

The full PRECISE-1K compendium, including the samples for this study, was used to compute iModulons using our previously described method ^41,77^. The log(TPM) dataset **X** was first centered such that wild-type *E. coli* MG1655 samples in M9 minimal media with glucose had expression values of 0 for all genes. Independent component analysis was performed using the Scikit-Learn (v0.19.0) implementation of FastICA ^78^. We performed 100 iterations of the algorithm across a range of dimensionalities, and for each dimensionality we pooled and clustered the components with DBSCAN to find robust components which appeared in more than 50 of the iterations. If the dimensionality parameter is too high, ICA will begin to return single gene components; if it is too low, the components will be too dense to represent biological signals. Therefore, we selected a dimensionality which was as high as possible without creating many single gene components, as described ^77^. At the optimal dimensionality, the total number of iModulons was 201. The output is composed of matrices **M** [genes x iModulons], which defines the relationship between each iModulon and each gene, and **A** [iModulons x samples], which contains the activity levels for each iModulon in each sample.

For each iModulon, a threshold must be drawn in the **M** matrix to determine which genes are members of each iModulon. These thresholds are based on the distribution of gene weights. The highest weighted genes were progressively removed until the remaining weights had a D’agostino K^2^ normality below 550. Thus, the iModulon member genes are outliers from an otherwise normal distribution. iModulon annotation and curation was performed by comparing them against the known TRN from RegulonDB ^79^. Names, descriptions, and statistics for each iModulon are available from the PRECISE-1K manuscript ^41^, iModulonDB ^20^, and **Table S2.**

#### Differential iModulon activity analysis

DiMAs were calculated as previously described ^10,21^. For each iModulon, a null distribution was generated by calculating the absolute difference between each pair of biological replicates and fitting a log-normal distribution to them. For the groups being compared, their mean difference for each iModulon was compared to that iModulon’s null distribution to obtain a p-value. The set of p-values for all iModulons was then false discovery rate (FDR) corrected to generate q-values. Activities were considered significant if they passed an absolute difference threshold of 5 and an FDR of 0.1. The main comparison in this study was between the starting strain at 250 μM PQ (n = 2) and the combined set of all evolved strains at 250 and 750 μM PQ (n = 61). Performing the comparison using both concentrations of PQ ensures that our comparison captures all of the major effects of tolerization. The set of DiMAs was similar when performing the comparison at just one or the other concentration.

We also performed a brief DEG analysis, which used the same algorithm as above but with individual gene expression values instead of iModulon activities.

#### iModulon explained variance calculation

The explained variance for each iModulon in this study was calculated using our workflow ^21^. Since iModulons are built on a matrix decomposition, the contribution of each one to the overall expression dataset can be calculated. For each iModulon, the column of **M** and the row of **A** for the evolved samples in this study were multiplied together, and the explained variance between the result and the full expression dataset was computed. These explained variance scores were used to size the subsets of the treemap in **Figure 1H.** Note that the variance explained by ICA is ‘knowledge-based’ in contrast to the ‘statistic-based’ variance explanation provided by the commonly used principal component analysis (PCA).

#### ME modeling

We used OxidizeME, a genome-scale model of metabolism and expression (ME) with ROS damage responses ^3^. Models used for flux maps were constrained using phenotypic data (glucose uptake rate and growth rate) and expression data as previously described ^24,25^. In order to force PQ cycling in the model, the lower bounds for the ‘PQ2RED_FWD_FLAVONADPREDUCT-MONOMER_mod_fad’ and ‘PQ1OX_FWD_SPONT’ were set to the same non-zero value and iterated over. Additionally, the former reaction was amended to accept NADH as an electron donor by editing the stoichiometry. PQ cycling sweeping calculations were performed by sampling various lower bounds to identify the range the model could support growth, and then sweeping 100 uniform values within that range. The total NADH produced through the TCA cycle was calculated by summing the fluxes for the ‘MDH’ and ‘AKGDH’ metabolic reactions. The percentage of the proteome allocated to the TCA cycle was calculated using the solutions from each model, specifically the translation fluxes:

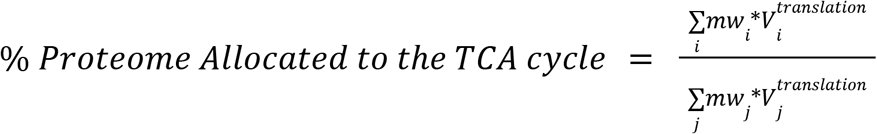

Where *mw_i_* and 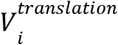 represents the molecular weight and translation flux of the *i*th protein in the TCA cycle, and *mw_j_* and 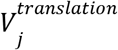 represents the molecular weight and translation flux of the jth protein the entire model. The damaged portion of the proteome was calculated as follows:

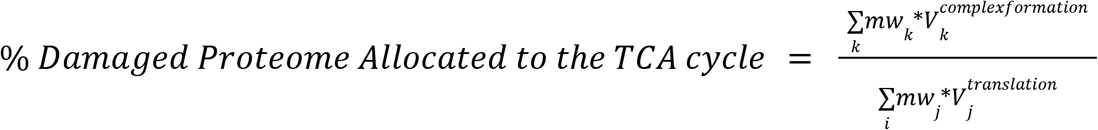

Where *mw_j_* and 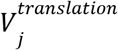 are the same variables above, and *mw_k_* and 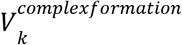 correspond to the kth protein in the table below:

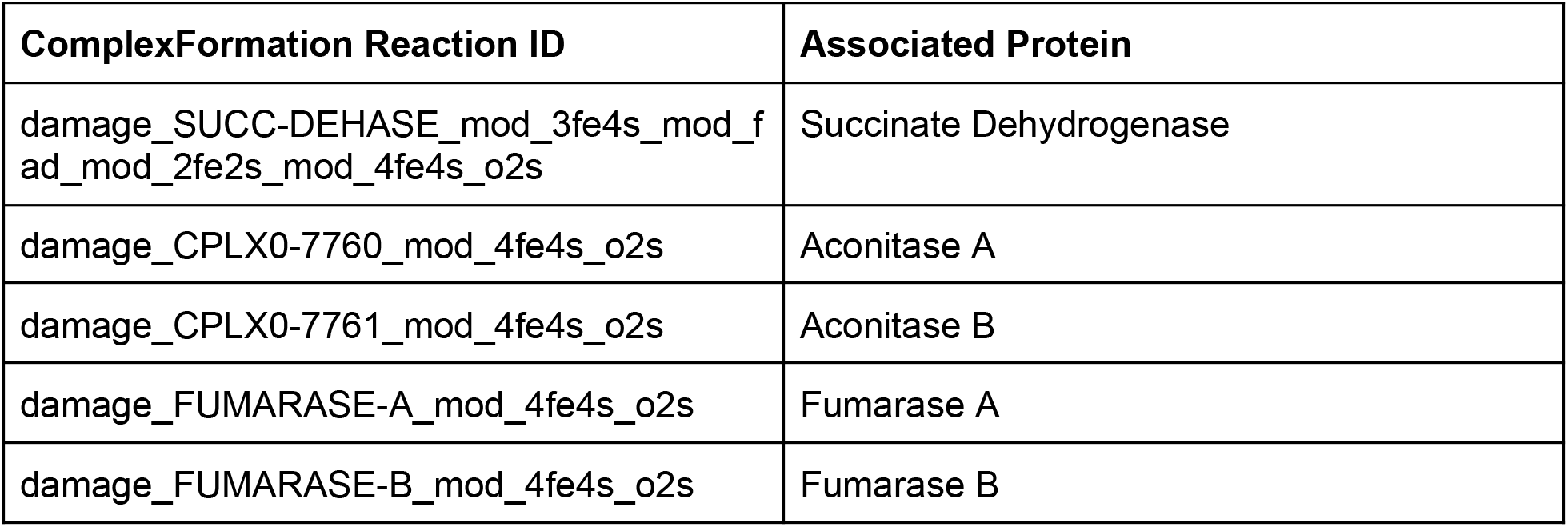

The undamaged portion of the proteome allocated to the TCA cycle was calculated as the difference between the total proteome allocated and the damaged proteome allocated.

### Additional resources

*iModulonDB: https://imodulondb.org/dataset.html?organism=e_coli&dataset=precise1k ALEdb: http://aledb.org/stats/?ale_experiment_id=1540*

## Supplemental Information

### Supplemental Figures

**Figure S1.**
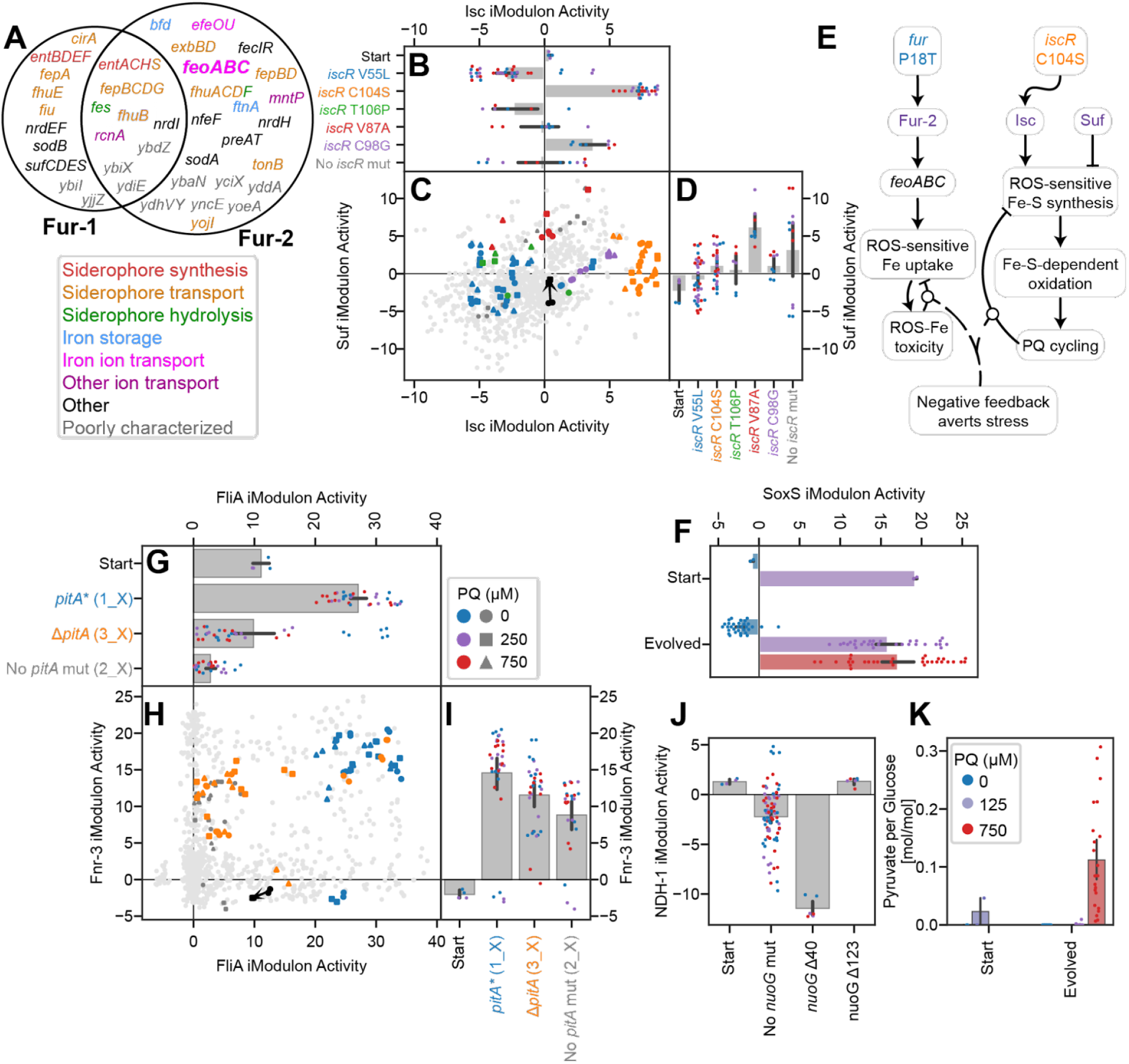
Additional insights from mutational, iModulon, and metabolic analysis. Bars indicate mean ± 95% confidence interval. **(A)** Venn diagram of the Fur-1 and Fur-2 iModulon genes, color coded by function. Ion transport and storage systems, which may be advantageous under ROS conditions, are enriched in Fur-2. **(B-D, H-J)** Scatter plots of iModulon activities with bar plots sharing axes. Light gray dots indicate other samples from PRECISE-1K. In (C) and (H), samples are colored by relevant mutations, and shapes indicate PQ concentrations according to the legends. A black arrow connects the starting strain samples between 0 and 250 μM PQ. In bar plots, point colors indicate PQ concentrations and label colors match with the scatter plots. (**B-D**) Suf and Isc iModulon activities, which are both regulated by IscR and encode distinct Fe-S cluster synthesis mechanisms (Suf is more robust to stress compared to Isc). (**E**) Knowledge graph linking two key TF mutations through their iModulons to negative feedback which averts stress. **(F)** SoxS iModulon activity is correlated with PQ in both starting and evolved strains (Pearson R = 0.72, p = 5.5*10^-15^). **(G-I)** FliA and Fnr-3iModulon activities by *pitA* mutation, showing an unexpected upregulation in the case of the frameshift pitA*, but not in the case of the pitA deletion. **(J**) NDH-1 iModulon activities. The NDH-1 iModulon consists of genes (nuoGHIJKLMN) that are controlled by ArcA and Fnr and are all downstream of the *nuoG* Δ40 mutation, which may create a terminator sequence. **(K)** Pyruvate production rates from exometabolomic characterizations of evolved strains. Note that the starting strain was characterized at 0 and 125 μM PQ (due to no growth at higher PQ), whereas the evolved strains were characterized at 0, 250, and 750 μM PQ. Pyruvate is secreted at high PQ levels, particularly by evolved strains which have downregulated PDH and the TCA cycle.

**Figure S2.**
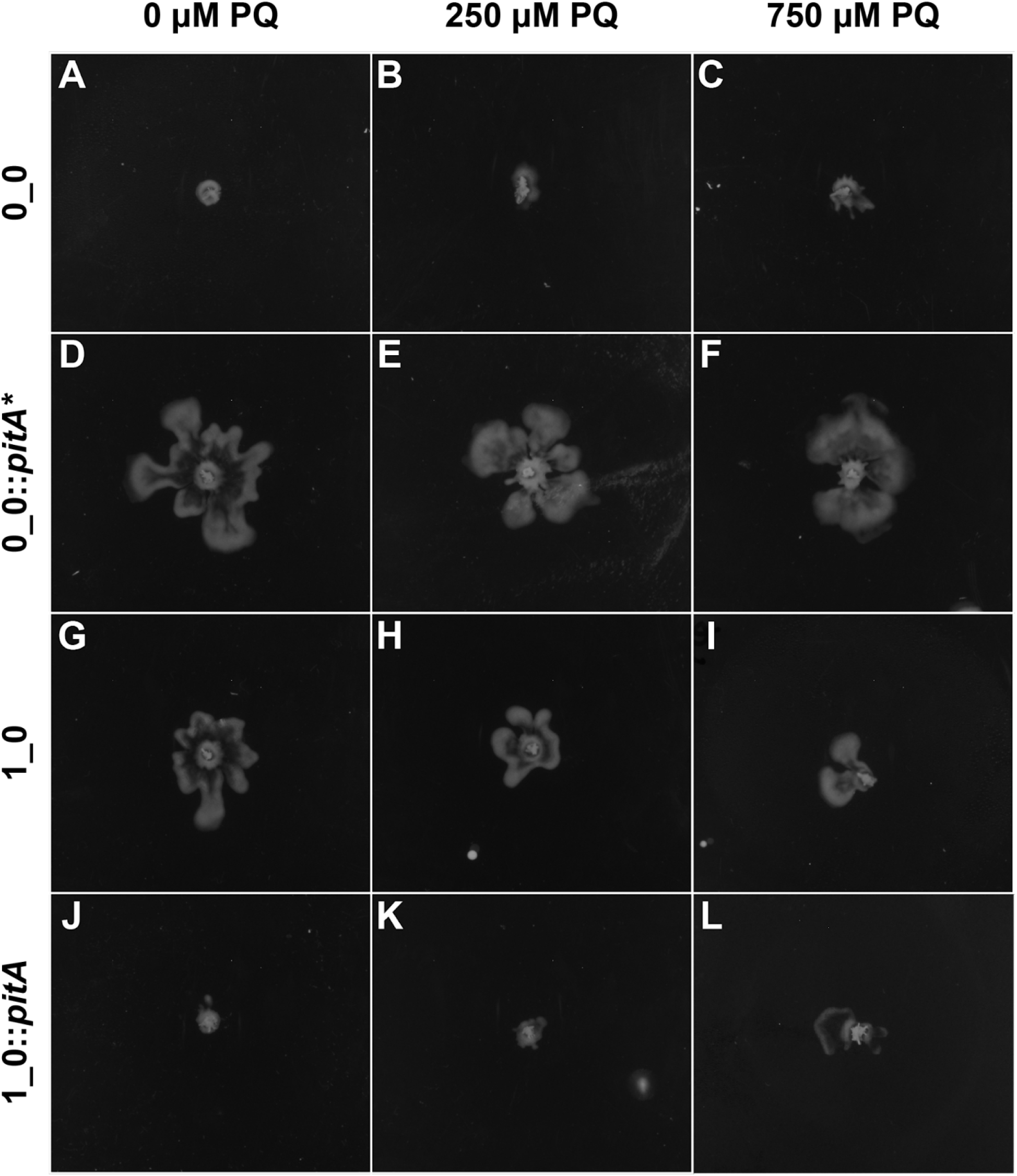
Swarming assays of pitA mutants. Cells were plated on agar in tryptone broth with glucose and the PQ concentration shown in the column headers. They were allowed to swarm for one day prior to image capture. The *pitA* mutant strains 0_0::*pitA\** (**D-F**) and 1_0 (**G-I**) swarmed, while wild type *pitA* strains (**A-C; J-L**f did not. Panels **A** and **D** are shown in **Figure 4K**

**Figure S3.**
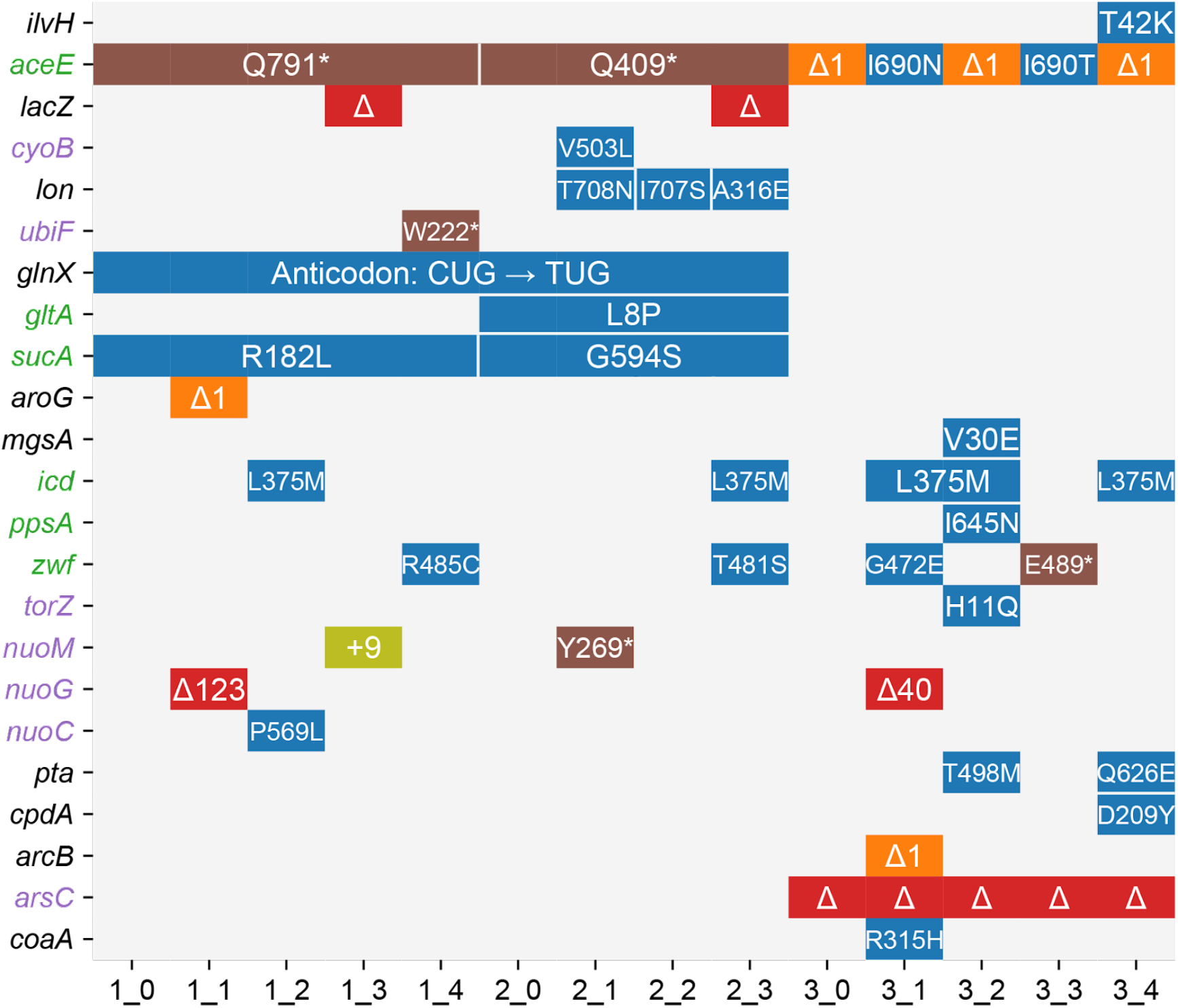
Mutations in genes relevant to metabolism. Colored blocks share a mutation in the given strain and gene (blue: missense SNP; brown: nonsense SNP; orange: frameshift deletion less than 3 bp; red: large deletion affecting gene; olive: insertion that does not cause a frameshift). Gene names are colored by type (green: central carbon metabolic enzyme; purple: redox enzyme; black: other gene relevant to metabolism). Silent mutations, the e14 deletion, and promoter mutations are omitted.

**Figure S4.**
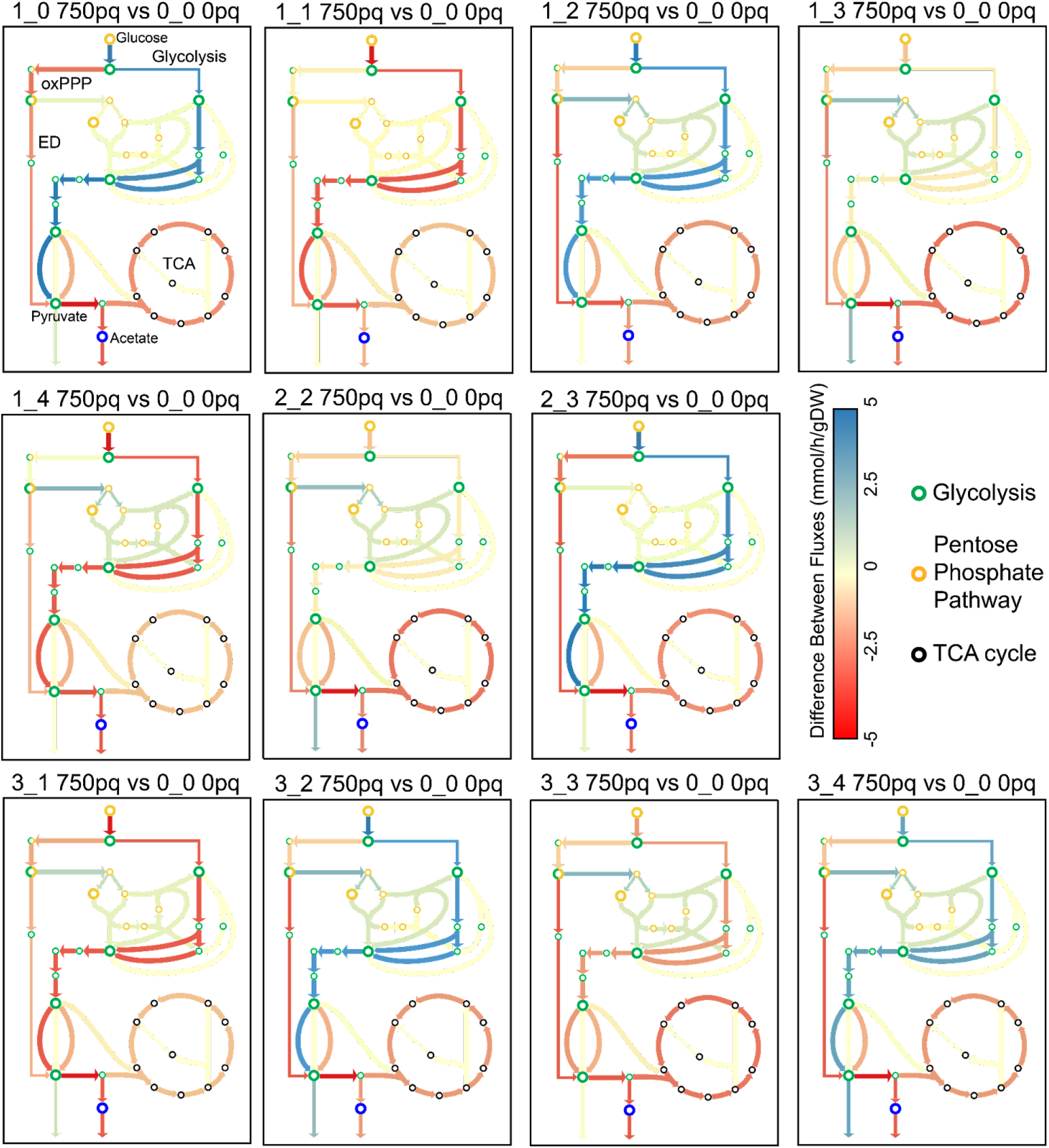
The constrained OxidizeME model predicts the flux distribution change in central metabolism after evolution. Flux distribution changes from specific OxidizeME models, constrained by RNAseq, growth, and glucose uptake data. TCA cycle flux always decreases after evolution (**Figure 6D**), and glycolytic flux varies with glucose uptake rate. Note that glucose uptake increases in evolved strains relative to the stressed starting strain, but some strains have more or less glucose uptake relative to the unstressed starting strain.

### Supplemental Tables

***Table S1. Mutations in ROS tolerized strains*.**

Mutations details, position, type, sequence change, and affected genes were generated by the ALEdb mutation calling pipeline ^8^. Hypothetical effect descriptions, both levels of categorization, and figure references were manually curated. ‘Treemap Category’ was used to generate **Figure 1D.** Columns labeled with strain numbers indicate presence or absence of the mutation in the given strain.

***Table S2. Significantly differentially activated iModulons in ROS tolerized strains*.**

List of significant DiMAs from the comparison shown in **Figure 1F**, in which evolved strains under 250 and 750 μM PQ were compared against the starting strain under 250 μM PQ. ‘Difference’ refers to the difference between the mean activity level of both groups, which has an absolute value greater than 5. ‘P-value’ is the false discovery rate corrected p-value for the statistical comparison, which is less than 0.1. ‘Explained Variance’ is the explained variance of the iModulon in the study samples, which was used with ‘Treemap Category’ to generate **Figure 1H.** ‘Treemap Category’, ‘Confidence’, ‘General Notes’, ‘Start Strain’, and ‘Evolved Strain’ were manually curated, with the latter two columns describing predicted regulatory mechanisms explaining the iModulon’s behavior in the respective samples. Remaining descriptive columns are copied from the PRECISE-1K curation of these iModulons ^41^. See iModulonDB.org for details of each iModulon, including its member genes, activity levels across over 1000 conditions including those from this study, and overlap with associated regulons.

### Note S1. PQ tolerant strains modify membrane transport, related to Figure 3

In the 3_0 strain and its subsequent evolutions, we do not observe the *emrE* amplification. However, our mutation caller predicted a 9-base pair (bp) insertion 39 bp upstream of *emrE* in these strains, consistent with IS1 insertions that can affect transcription or translation ^80^. We do not observe an iModulon signal in the transcriptome of these strains (**Figure 3C**). However, we do have evidence that increased expression of *emrE* provides an evolutionary benefit. Therefore, we hypothesize that this mutation would increase translation of EmrE.

The 3_0 strain and its descendants have a large deletion containing 26 genes (**Figure 3B**). The deletion may have been mediated by the *insH11* transposase at its 3’ end. Similarly to the *emrE* Amp iModulon discussed in the text, the Del-1 iModulon captured the effect of this change in the genome on the composition of the transcriptome. It showed a strong decrease in activity in the strains harboring the deletion (**Figure 3D**). The deleted segment contained a variety of genes, making it difficult to deduce its benefit to ROS tolerization. However, we note that it contained four transporter genes: *yhhJ, pitA, dtpB*, and *arsB*. Removal of one or several of these transporters may have decreased PQ influx or helped to prevent influx of other oxidized molecules that resulted from oxidative damage.

In addition to the transporters in the Del-1 iModulon, other deleted genes may have been important for the PQ tolerance of the 3_0 strain and its subsequent evolutions (**Figure 3B**). These include universal stress response regulators *uspBA*, reductases *gor, arsC*, and *yhiN*, or ribosome-related genes *rbbA, rsmJ*, and *rlmJ. yhhJ* and *yhiN* are uncharacterized genes with putative assignments, and these results support their potential role in PQ stress.

The *oppABCDF* operon was a common target of mutations. Nine of the eleven second-generation strains acquired the same 1,199 bp deletion of the *insH21* IS5 element upstream of it, and one strain, 1_1, deleted the entire operon and its surrounding genes. The deletion was captured by an iModulon (Del-2). The activity of this iModulon shows a downregulation in the deleted strain, and little change between the evolved strains with and without the upstream deletion (**Figure 3E**). Since *oppABCDF* is known to be a promiscuous tripeptide transporter that prefers positively charged substrates ^81^, it should be considered as a possible route of entry for PQ. The prevalence of the upstream deletion suggests that such a deletion provides improved tolerance, and there is an apparent benefit to a complete deletion of the entire operon. This leads us to predict that the upstream deletion negatively impacts *oppABCDF* translation, as has been suggested in past studies ^82,83^.

In addition to the genome-transcriptome-phenotype associations we analyze in depth, mutations on their own can predict putative new functions for their target genes. Therefore, we include all transporters mutated in this study in **Figure 3F** so that further research can explore their affinities for PQ and other oxidized compounds, as well as the effects of the observed SNPs.

### Note S2. Fur iModulon activities are variable and depend on *fur* mutations, related to Figure 4 and S1A

Fur, the ferric uptake regulator, regulates two main iModulons whose activities have a non-linear activity relationship which has been described in detail previously^46^ (**Figure 4B-D**). Fur-1 mostly contains genes for siderophore synthesis and transport (**Figure S1A**) which are derepressed under more extreme iron starvation conditions. Fur-2 contains ferrous iron transport genes, as well as siderophore transport and hydrolysis systems, which are derepressed more easily under relatively higher iron concentrations. The activities of the two iModulons form a logarithmic curve (**Figure 4C**), which captures the nonlinear effect of Fur on the composition of the transcriptome.

ROS demetallates iron enzymes and oxidizes iron(II) to iron(III) ^3,84^. Thus, PQ would induce higher intracellular iron concentrations that could be sensed by Fur and cause repression of both iModulons (black arrow, **Figure 4C**) ^85^. This hypothesis is consistent with the starting strain’s behavior. After evolution, a decrease in oxidative stress leads to a general upregulation of the Fur-1 and Fur-2 iModulons (p = 0.031 and 0.034, respectively).

The evolved strains exhibit a great degree of variation along the Fur curve (**Figure 4C**). Since many different factors could perturb iron concentrations for each culture (e.g. local ROS concentrations, trace element mixture variability, enzyme metallation levels, etc.), and Fur is highly sensitive to those concentrations, we believe that this variation is to be expected.

The mutation *fur* P18T was observed in three separate strains (1_2, 1_4, and 3_4). Strains with this mutation tend to be above the trend line in the Fur scatterplot (**Figure 4E**), suggesting a higher preference for expressing Fur-2 relative to Fur-1. The strains with this mutation specifically upregulated the *feoABC* genes, which are members of Fur-2 (**Figure 4F, S1A**).This transporter system may be highly beneficial under ROS conditions because it directly couples demetallation of an Fe-S cluster to iron transport, allowing for rapid decreases in iron acquisition when ROS levels are high ^47^.

Two other mutations were also observed in *fur*. H71Y in 1_3 tends to decrease expression of both iModulons, perhaps by strengthening Fur binding. This would potentially have the benefit of preventing iron toxicity. However, this strategy was not utilized by any other strains and it may have also hampered iron homeostasis in situations where local iron concentrations are low. The other mutation, A53G in 3_2, did not have a detectable effect on the transcriptome.

### Note S3. IscR mutations modify the balance between Fe-S cluster synthesis mechanisms and wildtype SoxS ensures ROS readiness, related to Figure 4 and S1F

IscR regulates two separate iron-sulfur (Fe-S) cluster synthesis systems which have iModulons, Isc and Suf^86^. Isc is associated with housekeeping Fe-S synthesis, whereas Suf is robust to iron starvation and ROS stress ^87–90^. Across our strains, we observed 5 mutations in *iscR*, and each associated with a particular region in a scatter plot of Suf and Isc iModulon activities (**Figure S1B-D**). Interestingly, most mutations do not strongly upregulate the ROS-tolerant Suf system (**Figure S1D**), and they either increase or decrease the expression of the Isc system (**Figure S1B**).

The particular regions in **Figure S1C** that were selected by the strains are somewhat unexpected. *iscR* C104S has been previously reported ^3,91^. The mutation is in IscR’s own Fe-S binding site, which causes it to maintain an unbound state that should de-repress Isc and activate Suf ^90,91^. We observe a strong upregulation of Isc in these strains, with more modest increases in Suf iModulon activity. The other most common mutation, *iscR* V55L, seems to downregulate Isc while also keeping Suf near basal levels. Given that ROS stress induces Fe-S cluster damage and Suf is significantly better at handling ROS stress ^89^, we would initially expect mutations which upregulate Suf to be more effective under the ALE conditions and therefore be enriched in these strains. We only see one mutation, *iscR* V87A, which seems to achieve that.

One possible explanation for this unexpected outcome is that the proteomic cost of the systems, particularly Suf, selects against strains which allocate too many resources towards Fe-S synthesis; this explanation has been modeled in a ME flux balance analysis ^3^. However, another possibility relates to the control of electron flux described in the metabolic section of the text: many redox enzymes, including some in respiration and the TCA cycle, contain Fe-S clusters ^92^. Damage to these enzymes by high ROS slows oxidative metabolism. This would charge fewer electron carriers and therefore slow the PQ cycle, allowing the cell to recover. It would therefore be better to express less Suf so that Fe-S synthesis would remain sensitive to ROS – using Isc or less of both systems would strengthen the coupling between ROS and respiration as a means of controlling the PQ cycle (**Figure 4K**). Thus, like the Fur P18T mutation (**Note S2**), this mutation enables a negative feedback loop, which aids in slowing oxidative metabolism and PQ cycling when stress is high (**Figure S1E**). This agrees with metabolic insights discussed in **Figures 6** and **7**.

While discussing Fe-S clusters, it is also worth noting that every strain mutated the putative Fe-S cluster repair gene *ygfZ*. This provides evidence for its role in Fe-S cluster homeostasis and motivates further study (**Table S1**).

Interestingly, there was a lack of mutations affecting *soxS*, the regulator of processes that remove the ROS superoxide ^93^. SoxS iModulon activity is highly correlated with PQ in the starting and all evolved strains (**Figure S1F**; Pearson R = 0.72, p = 5.5*10^-15^). The lack of mutations suggests that ROS readiness is preserved by using wild-type *soxS*.

### Note S4. PitA mutants’ upregulation of motility also upregulates anaerobic metabolism, providing a benefit to the strains, related to Figure 4 and S1

We propose the following possible mechanism for the benefit that motility upregulation provided to the *pitA* mutants (**Figure 4H-K**). The gene *aer*, which is upregulated as part of the FliA iModulon, mediates aerotaxis and would therefore allow cells to swim away from locally high concentrations of ROS ^94,95^. However, in our well-mixed cultures, there may not be local high ROS concentrations. In addition to its role in chemotaxis, *aer* helps to upregulate the Entner-Doudoroff pathway and anaerobic metabolism ^49^, a tendency which can be observed in the iModulon activities of our strains. Each of the anaerobic iModulons, Fnr-3 in particular, is slightly upregulated by the strains with *pitA\** (**Figure S1G-I**). An increase in anaerobic metabolism would help to prevent PQ cycling as described in the text. Thus, a decrease in oxidative metabolism is also achieved by the cells through this very non-conventional mechanism. An added benefit may lie in the expression of *fliZ*, a member of the FliA iModulon, which is known to antagonize RpoS and would therefore be expected to promote growth ^96^.

### Note S5. iModulons related to specific metabolites reflect decreased stress after tolerization, related to Figure 5

The ppGpp iModulon contains a large set of growth-related genes regulated by the master regulator ppGpp ^97^. It follows a similar pattern to the Translation iModulon, suggesting that ppGpp concentrations decline after evolution (**Figure 5C**).In addition to the Translation and ppGpp iModulons, a few other differentially activated iModulons with more specific functions are also likely to be responding to ppGpp levels, including the Nucleotide Stress, Glutarate, Efflux Pump, and Biofilm iModulons (**Table S2**).

Ribose concentrations are sensed by RbsR ^98^, which represses the Ribose iModulon in its presence. Ribose is produced as part of the pentose phosphate pathway (PPP), which is the primary pathway for producing NADPH to detoxify ROS. Upon initiation of oxidative stress, PPP flux increases, producing ribose ^99^. Oxidative stress also slows growth and DNA synthesis, which will decrease ribose utilization. We therefore expect an increase in ribose concentrations in the starting strain upon PQ stress, which is observed as a decrease in Ribose iModulon activity (**Figure 5H**). In the evolved strains, flux shifts towards glycolysis and away from the PPP, producing less ribose. They also synthesize more DNA to support faster growth, using ribose. Therefore, Ribose iModulon activity increases relative to the starting strain, while still exhibiting a negative correlation with PQ.

The Purine iModulon is regulated by PurR and ppGpp, and its activation pattern in our samples (**Figure 5I**) mirrors that of the Translation and ppGpp iModulons (**Figure 5E**). This activation may be explained by direct action by ppGpp, or via PurR, which represses these genes in the presence of hypoxanthine or guanine ^100^. The faster growing evolved strains would perform more DNA replication and RNA synthesis, and therefore require purine synthesis, depleting the metabolites which are sensed by PurR and de-repressing the iModulon.

Changes in Cysteine-1 iModulon activities may be explained by increased ROS readiness and subsequent improvement in amino acid homeostasis (**Figure 5J**). This iModulon is regulated by CysB, which can be inhibited by cystine and other oxidized sulfur compounds ^101,102^. Cysteine is very easily oxidized ^103,104^, which may explain the dramatic downregulation of the iModulon upon PQ addition in the starting strain. The evolved strains with PQ have significantly higher Cysteine-1 activity compared to the parent strain with PQ, due to the success of their tolerization strategies.

The Copper iModulon, which contains copper efflux genes regulated by CueR, CusR, and HprR, is downregulated in the evolved strains (**Figure 5K**). Copper is redox-sensitive, and its efflux depends on the proton-motive force (PMF) or ATP ^105^. It is also an important cofactor for various enzymes, including the superoxide dismutase *sodC* ^106^. Oxidative damage should decrease the PMF and ATP concentrations and alter the copper redox state, which would explain the iModulon’s upregulation in the stressed starting strain. The evolved strains downregulate this iModulon, reflecting improvements in metal homeostasis resulting from ROS tolerization.

The Arginine iModulon is generally upregulated in the evolved strains relative to the stressed starting strain. This set of genes is regulated by ArgR, which represses them in the presence of arginine, and is also influenced by ppGpp ^107^. The iModulon activity in the starting strain indicates that oxidative stress increases arginine concentrations. This activation may be due to a variety of reasons, including damage to polyamine synthesis pathways that use arginine as a precursor ^108,109^. Since the stressed evolved strains behave more like the unstressed starting strain, it appears that arginine homeostasis is restored by ROS tolerization.

### Note S6. Synergistic mutations in PDH and a tRNA balance the tradeoff, related to Figure 5

The growth/stress tradeoff is embodied by interactions between two mutations, which both occurred in both the 1_0 and 2_0 strains. First, *aceE* acquired a C→T nonsense SNP, creating an amber stop codon ^110^: Q791* in 1_0 and Q409* in 2_0. This mutation inactivated PDH and likely significantly decreased flux into the TCA cycle. While effective early in the evolution at decreasing PQ cycling, the change was extremely damaging. Interestingly, both 1_0 and 2_0 later acquired the same C→T SNP in the anticodon of the glutamine tRNA *glnX* ^111^. This second change enabled the mutant *glnX* to read through the initial *aceE* truncation, allowing for some functional PDH to be translated and utilized for energy generation. Due to competition between stop codon release factors and *glnX*, functional *aceE* translation would not return to wild type levels ^112^, but rather find an intermediate level which balanced the tradeoff.

We quantified the above relationship using ribosome profiling (**Figure 5C**). By measuring the fraction of ribosomes bound to the sequence before and after the truncating SNP, we demonstrated the near complete deactivation of *aceE* translation in the midpoint strain. In the 2_0 strain with both the *aceE* and *glnX* mutations, translation was partially restored (to a ratio of 0.23±0.08). Thus, synergy between these two mutations brokered a compromise between the energy and stress-generating effects of TCA cycle flux.

The 3_0 strain acquired a frameshift 1 bp deletion in *aceE* instead of the nonsense SNP. This meant that it could not employ a similar strategy to 1_0 and 2_0. However, two of its second generation derivative strains (3_1 and 3_3) had insertions at or near the deletion (**Figure S3**), which may have served a similar purpose in re-increasing PDH levels.

### Note S7. NADH dehydrogenase and other reductases may be PQ diaphorases, related to Figures 6 and S2

In addition to the NADH production-related mutations described in the text, we also observe NADH utilization-related mutations. Five strains acquired unique mutations in *nuoC, nuoG*, and *nuoM* of the NADH dehydrogenase complex (NDH-1). A 40 bp deletion within *nuoG* appears to induce early termination of transcription, since genes downstream of it are captured by the NDH-1 iModulon and strongly downregulated in the strain with the deletion (**Figure S1J**). Note that another strain deleted 123 bp in a nearby region of the same gene, but we do not observe early termination in that strain. The prevalence of these mutations suggests a benefit to NDH-1 LOF under PQ conditions.

Cellular enzymes which catalyze PQ reduction are called PQ diaphorases, and three have been identified in *E. coli* by past studies ^61,113^. Those studies suggested that NADPH plays a larger role than NADH, but our mutations preferentially affect NADH production and NDH-1. It is possible that transhydrogenases first convert NADH to NADPH^114^ prior to the PQ cycle. Alternatively, NDH-1 and other mutated NADH reductases from this study (e.g. *cyoB, ubiF, torZ*, and *irxC;* **Figure S3**) ought to be considered as potential PQ diaphorases. Though NDH-1 has not been implicated in PQ cycling in *E. coli*, this phenomenon has been observed in mammals ^60,62^.

### Note S8. Exometabolomics revealed secretion of pyruvate, consistent with TCA LOF, related to Figures 6 and S2

In addition to the glucose uptake rates discussed in the text, our exometabolomic physiological characterization quantified production of organic acids. We do not report the specific rates in the text because of the low signal to noise ratio for the low concentrations of these compounds in the early exponential phase region. However, it was interesting to note that pyruvate was secreted by several evolved strains at 750 μM PQ (**Figure S1K**). This is consistent with the expected decreased function of PDH and TCA that was predicted from the mutations, and with the intracellular pyruvate concentration increase predicted for all PQ levels in the evolved strains (**Figure 7E, 7H; Note S9**). We also observed acetate production by all strains.

### Note S9. iModulon activities shift tolerant strains towards anaerobic metabolism and glycolysis, related to Figure 7

ArcA is part of the ArcAB two-component system, which senses the ratio of reduced to oxidized quinones in the ETC ^64^. In the starting strain, oxidative stress from PQ shifts this ratio toward oxidation, causing ArcAB to be less active and derepress the ArcA iModulon. As strains evolve, they experience less oxidative stress due to their transport and TCA cycle mutations. This lowered stress leads to a more reduced quinone pool, an increase in ArcAB activity, and repression of the ArcA iModulon (**Figure 7A**). The ArcA iModulon contains aerobic growth genes such as oxidoreductases and cytochromes, so its repression will encourage anaerobic metabolism, fermentation, and a decreased reliance on NADH.

There are two strains which have mutations in the ArcAB two-component system, affecting ArcA iModulon activity. A frameshift in the sensor kinase *arcB* in 3_1 has a moderate derepressing effect, and an early stop in *arcA* in 1_1 had a stronger derepressing effect (**Figure 7A**). These two strains are an exception which appear to have struck a different balance in the growth/stress generation tradeoff compared to the other evolved strains. They express aerobic metabolism genes as well as the Fnr-activated anaerobic fermentation genes, which would enable them to use more energy producing pathways but could also exacerbate stress generation.

Fnr senses oxygen levels via oxidative damage to its Fe-S cluster and activates anaerobic metabolism genes when the cluster is intact ^65,115^. Its regulon is captured by three iModulons, whose activities behave similarly in this study (**Figure 7B-D**). The decrease in oxidative stress, as well as the success of iron-related mutations, help to maintain more active Fnr and therefore upregulate this iModulon.

The Cra iModulon captures a set of genes of glycolysis and carbohydrate catabolism genes which are repressed by Cra ^116,117^. Cra regulates these genes by acting as a flux sensor for glycolysis, since their suppression is activated by fructose-1,6-bisphosphate ^118^. We observe an increase in Cra iModulon activity in the evolved strains (**Figure 7F**), which both indicates and positively regulates an increase in glycolytic flux.

The Crp-2 iModulon contains mostly phosphotransfer (PTS) system genes which are activated by the master regulator Crp ^119^. Crp responds to cAMP levels in a biphasic manner, and cAMP levels themselves have complex regulation^120^. We observe a strong downregulation of the Crp-2 iModulon in the stressed starting strain, but a return to unstressed or intermediate levels in the evolved strains (**Figure 7G**). This change is consistent with a return to homeostasis, and may indicate a more active PTS, higher glucose uptake, and increase in ATP concentrations after evolution.

The LOF mutations in PDH and the TCA cycle should increase intracellular pyruvate concentrations, since pyruvate is the initial substrate for those reactions. The Pyruvate-2 iModulon is regulated by PyrR, which can sense pyruvate concentrations ^121^. Pyruvate-2 activity increases in the evolved strains (**Figure 7H**), which is consistent with this prediction. We also observe pyruvate secretion at high PQ levels (**Figure S1K; Note S8**), probably due to the oxidative damage to PDH and the TCA cycle causing so much pyruvate accumulation that it must be secreted.

## Notes

https://imodulondb.org/dataset.html?organism=e_coli&dataset=precise1k

https://aledb.org/stats/?ale_experiment_id=1540

https://www.ncbi.nlm.nih.gov/bioproject/PRJNA913892

### Competing Interest Statement

The authors have declared no competing interest.

